# Molecular and Structural Basis of Olfactory Sensory Neuron Coalescence by Kirrel Receptors

**DOI:** 10.1101/2021.03.09.434456

**Authors:** Jing Wang, Neelima Vaddadi, Joseph S. Pak, Yeonwoo Park, Sabrina Quilez, Christina A. Roman, Emilie Dumontier, Joseph W. Thornton, Jean-François Cloutier, Engin Özkan

**Author notes:** Correspondence (E.Ö.), (J.F.C.). These authors contributed equally (Jing Wang, Neelima Vaddadi, Joseph S. Pak).

## Abstract

Projections from sensory neurons of olfactory systems coalesce into glomeruli in the brain. The Kirrel receptors are believed to homodimerize via their ectodomains and help separate sensory neuron axons into Kirrel2- or Kirrel3-expressing glomeruli. Here we present the crystal structures of homodimeric Kirrel receptors and show that the closely related Kirrel2 and Kirrel3 have evolved specific sets of polar and hydrophobic interactions, respectively, disallowing heterodimerization while preserving homodimerization, likely resulting in proper segregation and coalescence of Kirrel-expressing axons into glomeruli. We show that the dimerization interface at the N-terminal IG domains is necessary and sufficient to create homodimers, and fail to find evidence for a secondary interaction site in Kirrel ectodomains. Furthermore, we show that abolishing dimerization of Kirrel3 *in vivo* leads to improper formation of glomeruli in the mouse accessory olfactory bulb as observed in *Kirrel3*^*-/-*^ animals. Our results provide strong evidence for Kirrel3 homodimerization controlling axonal coalescence.

## INTRODUCTION

In the vertebrate olfactory system, olfactory sensory neurons (OSNs) expressing the same olfactory receptor project their axons to a limited set of glomeruli in the olfactory bulb (OB), which allows for the creation of a topographic odor map in the OB (Mombaerts et al., 1996). In the mouse olfactory system, each OSN expresses a single olfactory receptor, which drives the expression of a limited number of cell surface receptors (Serizawa et al., 2006). Specifically, the expression of the cell adhesion receptors Kirrel2 or Kirrel3 in OSN populations is dependent on the odorant-induced activity of the olfactory receptors, and their expression patterns are mostly complementary: Kirrel2 expressing OSNs and Kirrel3 expressing OSNs segregate into separate glomeruli (Serizawa et al., 2006).

The accessory olfactory system, which is found in most terrestrial vertebrate lineages and is responsible for the detection of chemosignals that guide social behavior, has a similar structure (Brignall and Cloutier, 2015; Dulac and Torello, 2003), where vomeronasal sensory neurons (VSNs) expressing the same vomeronasal receptor project their axons into a specific set of 6 to 30 glomeruli in the accessory olfactory bulb (AOB) (Belluscio et al., 1999; Del Punta et al., 2002; Rodriguez et al., 1999). The coalescence of VSN axons into glomeruli are also shown to be dependent on Kirrel expression, where VSN activity evoked by chemosignals reduces Kirrel2 expression and increases Kirrel3 expression (Prince et al., 2013). The two Kirrel paralogs show mostly complementary expression patterns in VSNs of the VNO and in their axons innervating glomeruli of the AOB. Ablation of Kirrel2 or Kirrel3 expression in mice leads to improper coalescence of VSN axons, resulting in fewer and larger glomeruli being formed in the posterior region of the AOB, while double knock-out mice no longer have recognizable glomeruli (Brignall et al., 2018; Prince et al., 2013).

Members of the Kirrel (Neph) family of cell surface receptors were first identified in fruit flies for controlling axonal pathfinding in the optic chiasm and programmed cell death in retinal cells (Ramos et al., 1993; Wolff and Ready, 1991). These molecules have also been recognized for their role in myoblast function during muscle development, and their involvement in building the blood filtration barrier in glomeruli in kidneys (Donoviel et al., 2001; Ruiz-Gómez et al., 2000). The *C. elegans* homolog, SYG-1, has been demonstrated to specify targeting of synapses (Shen and Bargmann, 2003). Outside its function in the olfactory system, mammalian Kirrel3 has been shown to be a synaptic adhesion molecule necessary for the formation of a group of synapses in the hippocampus (Martin et al., 2015; Taylor et al., 2020), and is associated with autism spectrum disorders and intellectual disability (Bhalla et al., 2008; De Rubeis et al., 2014; Taylor et al., 2020).

Here, we investigate the molecular basis of Kirrel2- and Kirrel3-mediated glomerulus formation in the AOB. We present the three-dimensional structures of Kirrel2 and Kirrel3 in homodimeric complexes that reveal why Kirrels do not heterodimerize, allowing for proper adhesion and segregation of subsets of sensory axons. We engineer mutations that perturb the homophilic interactions, abolish dimerization and Kirrel-mediated cell adhesion. Finally, we introduce a non-homodimerizing amino acid substitution into the mouse *Kirrel3* allele, which leads to alterations in the wiring of the accessory olfactory system that phenocopy defects observed in *Kirrel3*^-/-^ animals. These results indicate that Kirrel3 function in the control of glomeruli formation depends on ectodomain interactions mediated by Kirrel homodimerization via its first immunoglobulin domain.

## RESULTS

### Structural characterization of Kirrel2 and Kirrel3 D1 homodimers

It has been proposed that Kirrel2 and Kirrel3 regulate the coalescence of OSNs and VSNs into glomeruli by homophilic adhesion through their ectodomains (Prince et al., 2013; Serizawa et al., 2006). Vertebrate Kirrel proteins have been shown to form homodimers, but not heterodimers (Gerke et al., 2005; Serizawa et al., 2006), which may be important for axonal segregation into distinct glomeruli. Previously, we determined the crystal structure of mouse Kirrel1 (also known as Neph1) at 4 Å resolution covering the first two immunoglobulin domains, and showed that homodimerization occurs by an interaction between the first domains (D1) of each monomer (Özkan et al., 2014). Due to limited resolution, however, the Neph1 electron density was missing for most side chains and details of the interaction interface could not be documented. Vertebrate Kirrels are remarkably similar in their extracellular regions, containing five immunoglobulin domains (Figure 1A) with sequence identities of 53 to 56% for D1 among the three mouse Kirrels (Figure 1B).

**Figure 1.**
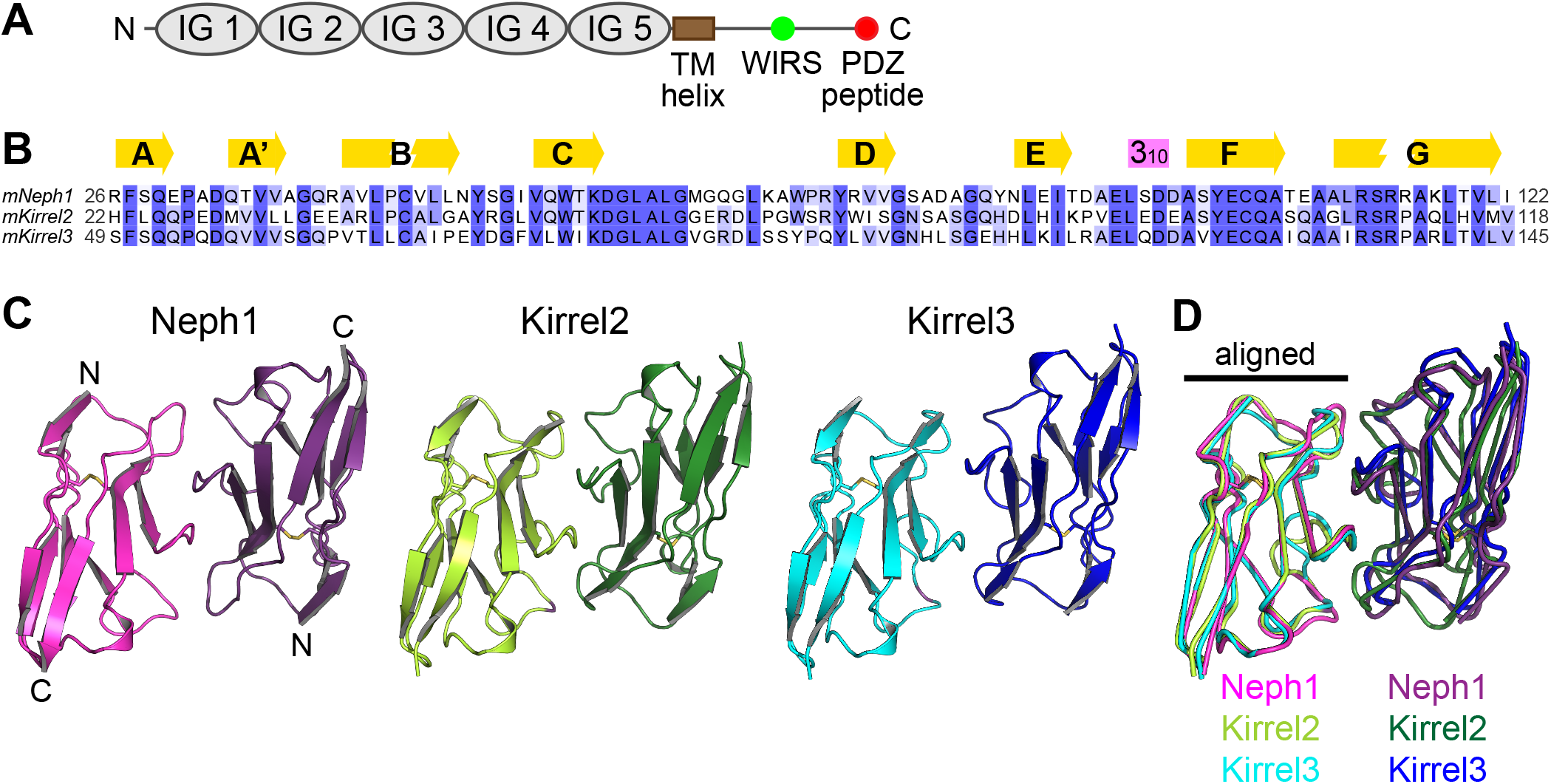
The three mouse Kirrel D1 homodimers share gross structural features. **A**. The conserved domain structure of Kirrels. **B**. A multisequence alignment of the mouse Kirrel D1 domains. Secondary structure elements, β-strands (yellow arrows) and a 310 helix (purple box) are shown above the alignment. **C**. The three mKirrel D1 dimers shown in cartoon representation. Neph1 homodimer structure is from Özkan et al. (2014; PDB ID: 4OFD). See Table S1 for structure determination statistics for Kirrel2 and Kirrel3 structures. **D**. The three mKirrel D1 homodimers are overlayed by aligning the chains shown on the left. Comparisons to structures of fly and worm Kirrel orthologs are provided in Figure S1.

In order to explain the molecular basis of homophilic interactions in the absence of heterophilic binding between the Kirrel paralogs, we determined the crystal structures of Kirrel2 and Kirrel3 D1 homodimers at 1.8 Å and 2.1 Å resolution, respectively (Figures 1C, Table S1). When superimposed, the Kirrel2 and Kirrel3 D1 domains have root-mean-square displacements (rmsd) of 1.2 Å to the Kirrel1 D1 domain, and align with an rmsd of 1.0 Å to each other (all Cα atoms). Comparison of the three Kirrel D1 homodimers show that Kirrels dimerize using the same interface and the dimeric architectures are shared (Figure 1D). The three D1 homodimers align with rmsd values of 1.3 to 1.6 Å. Similar dimers have been observed for homodimeric complexes formed by fruit fly Kirrel homologs, Rst and Kirre (Duf), and the heterodimeric SYG-1-SYG-2 complex, the *C. elegans* Kirrel homolog bound to Nephrin-like SYG-2 (Figure S1) (Özkan et al., 2014). This distinctive interaction architecture uses the curved *CFG* face of the IG fold to create dimers, and was recently identified to be common to a group of cell surface receptors involved in neuronal wiring, including Dprs, DIPs, IGLONs, Nectins and SynCAMs, as well as Kirrels and Nephrins (Cheng et al., 2019).

Out of 21 amino acids found in three structures to be at the dimer interface, ten are invariable among the three mouse Kirrels, mostly in the region following the *C* strand within the sequence KDGLALG, and the *F* and *G* strands. The conserved Gln128 (Kirrel3 numbering) in the *F* strand makes hydrogen bonds to the main chain of the conserved stretch in the *CD* loop in all three Kirrel structures (Figure S2). Beyond this similarity, there are stark differences in the Kirrel2 and Kirrel3 interfaces that likely underlie exclusive homodimerization. Kirrel2 has a hydrogen bonding network at the core of the interface, centered at Gln52 at the two-fold symmetry axis (Figure 2A), while the hydrogen bonding network is missing in Kirrel3, which has Leu79 at the symmetry axis instead (Figure 2B). The polar vs. hydrophobic interactions at the Kirrel2 and Kirrel3 interfaces, respectively, would preclude heterodimerization of the two Kirrels.

**Figure 2.**
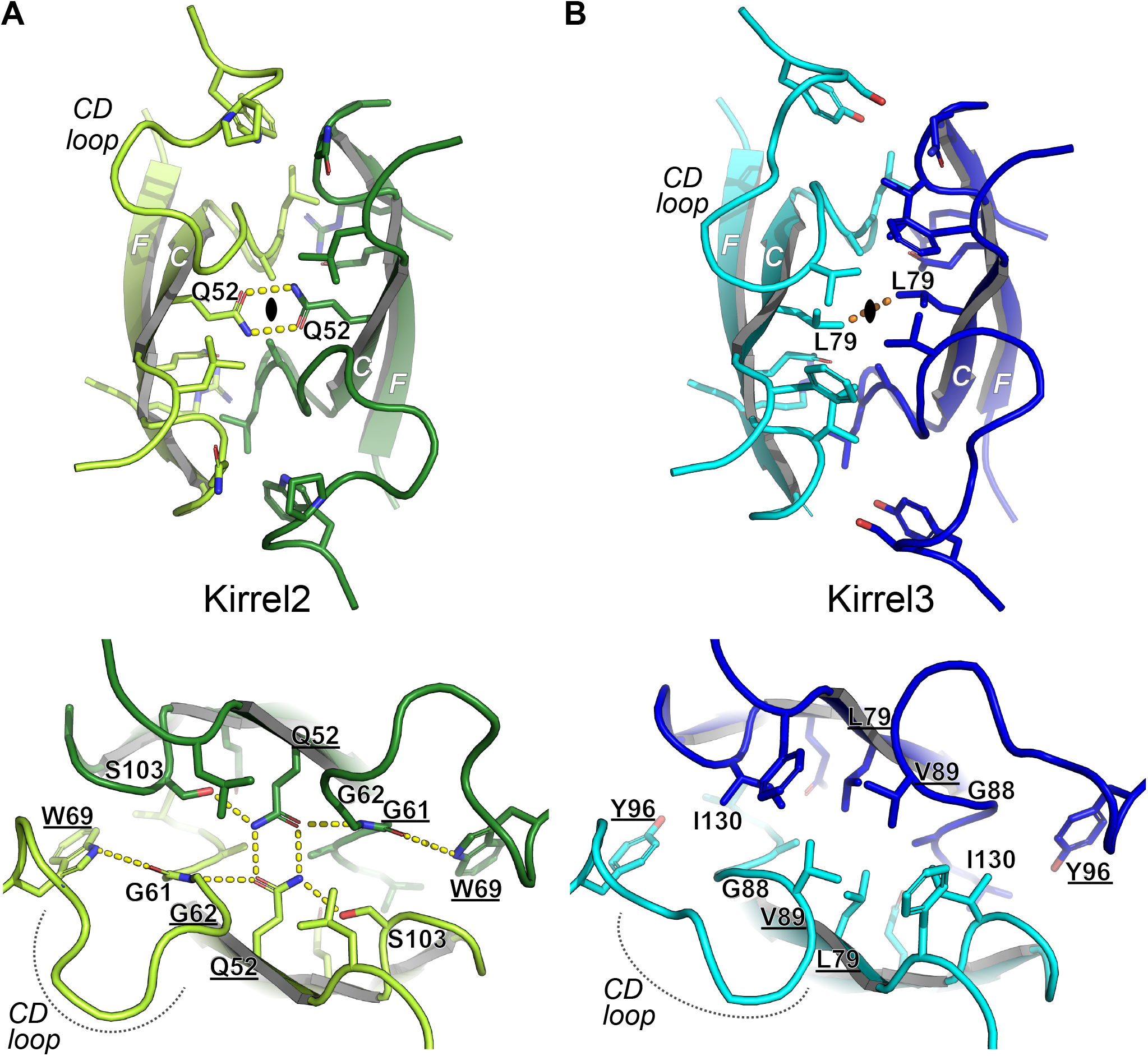
Kirrel2 and Kirrel3 have incompatible chemistries at their dimerization interfaces. **A**. Kirrel2 homodimerization interface. The closed oval represents the two-fold symmetry axis surrounded by the bidentate hydrogen bonding (yellow dashes) of two Q52 residues from Kirrel3 monomers. The interface includes an extensive set of hydrogen bonds. **B**. Kirrel3 homodimerization interface. None of the hydrogen bonds depicted in (A) are present in the Kirrel3 interface. The hydrogen bonds between the Q52 side chains at the Kirrel2 dimer symmetry axis are replaced by van der Waals contacts between the L69 side chains (orange dashes) in Kirrel3. The loop connecting the C and D strands are significantly different between the two Kirrel structures. The parts of the interface with conserved hydrogen bonding between Kirrel2 and Kirrel3 are shown in Figure S2. Underlined amino acids vary among ancestral vertebrate sequences (see Figure 3C) and are strong candidates to be specificity determinants.

### Evolutionary origins of vertebrate Kirrels and homodimeric specificity

To understand how Kirrel homodimeric specificities arose in vertebrates, we next analyzed the evolutionary histories of Kirrels. All major bilaterian taxa contain Kirrel homologs, which preserve the five-IG domain extracellular region and a cytoplasmic PDZ peptide. We created a maximum likelihood (ML) phylogeny for Kirrels using a multiple sequence alignment of 90 ectodomains (Figure 3A and Suppl. Data 1). The three vertebrate Kirrels arose as a result of gene duplications and are more closely related to each other than to invertebrate Kirrels. Kirrel duplication in invertebrates is rare, with one notable exception being *Drosophila*, which has two Kirrels (Rst and Kirre). After gene duplications within vertebrates, there have been several gene losses; this, combined with extensive sequence divergence between vertebrate and invertebrate Kirrels, makes it difficult to accurately reconstruct the branching order of vertebrate Kirrels. Nevertheless, we observe strong bootstrap values for cyclostome Kirrel1 forming a single clade with gnathostome Kirrel1 and excluding Kirrel2 and Kirrel3. This strongly implies that Kirrel2 and Kirrel3 arose from a gene duplication within gnathostomes and are more closely related to each other than to Kirrel1 in support of our ML tree.

**Figure 3.**
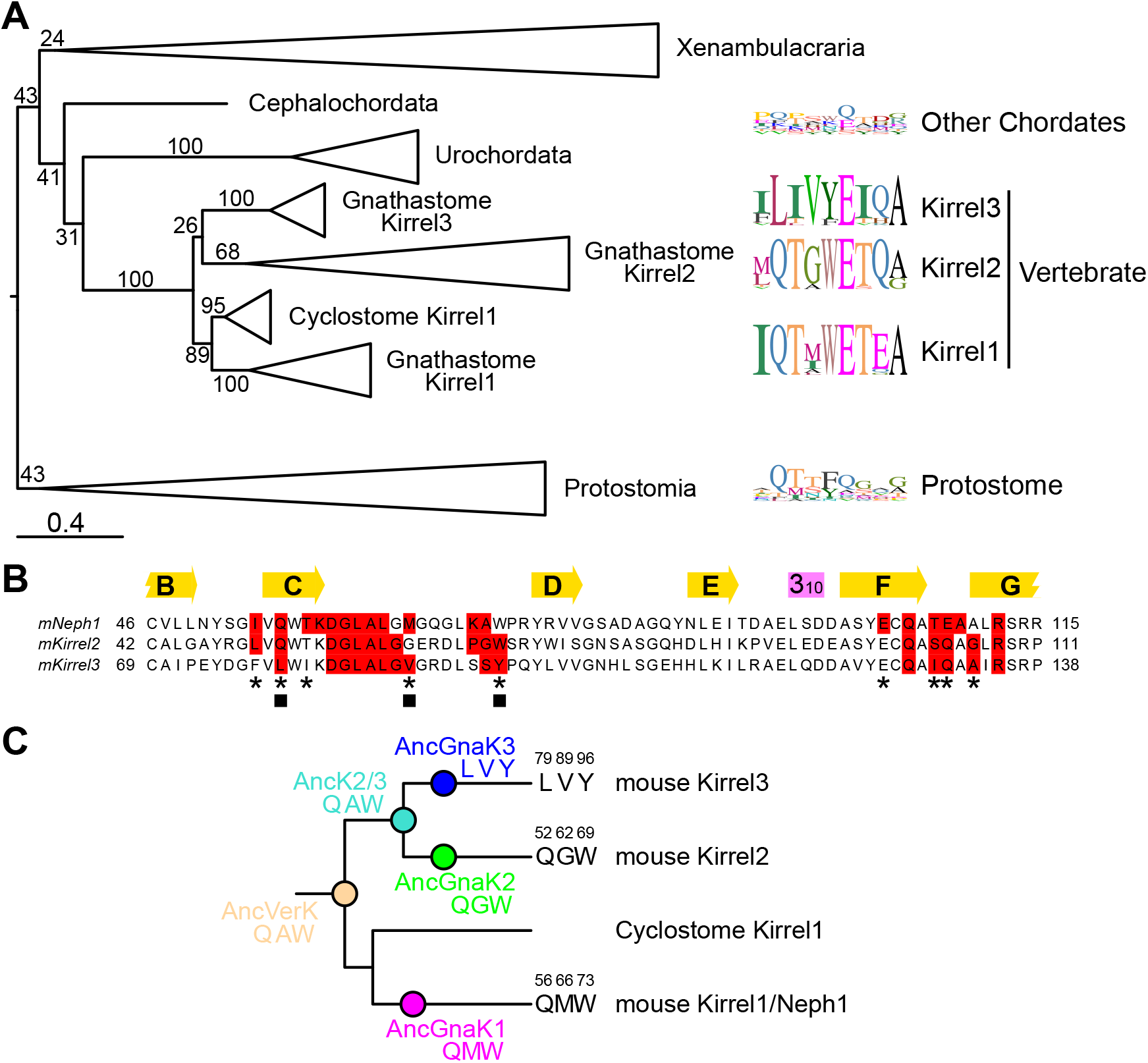
Phylogenetic analysis of Kirrels. **A**. Maximum likelihood tree for Kirrels. The scale bar below represents 0.4 substitutions per site. Numbers on the tree are bootstrap values supporting the adjacent node. The sequence logos to the right show the prevalence of amino acids in selected positions at the Kirrel dimerization interface for specific taxa, placed next to their branch in the phylogenetic tree. See Source Data 1 for the tree file with bootstrap support values. **B**. Sequence alignment showing all amino acids at the interface: red boxes; 4 Å cutoff used for identifying an interface amino acid. The selected residues used in sequence logos in (A) are marked with an asterisk below. Three positions at the interface that vary among ancestral sequences are labeled by closed squares. **C**. The three varying residues among sequence reconstructions of the three ancestral gnathostome Kirrels, the Kirrel2/3 ancestor and the ancestral vertebrate Kirrel are shown on the tree. These positions are underlined in the structural views of the interface in Figure 2. See Figure S3 for complete sequences of inferred ancestral D1 domains.

To understand the evolution of Kirrel interaction specificity, we mapped variable interface residues in separate Kirrel clades (Figures 3A and 3B). The sequence logos show that Kirrel1 and Kirrel2 are more similar at their dimerization interfaces, while Kirrel3 diverged and mutated key positions to non-polar amino acids. Surprisingly, much of the variability observed at these positions in vertebrate Kirrels are also seen in invertebrate Kirrels, suggesting that these sites are permissive for multiple modes and specificities of binding.

Next, we inferred ancestral sequences for Kirrel D1 domains in the vertebrate and gnathostome nodes on the phylogenetic tree (Figures 3C and S3). There are nine variable positions between the common Kirrel2/Kirrel3 ancestor and the ancestral gnathostome Kirrel3. Three of these positions are at the interface (labeled with closed squares in Figure 3B), including two that are part of the hydrogen bonding network seen in Kirrel2 (Q52 and W69), and one contributing strongly to the hydrophobic nature of the Kirrel3 interface (V89 in Kirrel3, G62 in Kirrel2) (see underlined amino acids in structures in Figure 2). Overall, these ancestral sequences support our structural observations that the gnathostome Kirrel3 homodimer interface gained non-polar interactions which resulted in loss of interactions with its Kirrel2 paralogs.

### Mutagenesis of the Kirrel2 and Kirrel3 D1 dimerization interfaces

To confirm that the Kirrel2 and Kirrel3 D1 interfaces observed in crystal structures mediate dimerization in solution, we set out to mutate these interfaces and test for loss of dimer formation. In size exclusion chromatography (SEC) experiments, Kirrel2 D1 and Kirrel3 D1+D2 proteins elute at a volume that roughly corresponds to that of their dimeric sizes (Figures 4A, 4B and S4A-F). This dimerization is concentration-specific; size-exclusion runs with diluted samples elute at later volumes, indicating a fast equilibrium between dimeric and monomeric forms of Kirrels. Interface mutations of mouse Kirrel2 D1 and Kirrel3 D1+D2 were tested for dimerization at various concentrations based on SEC elution volumes, which allowed us to rank these mutations for their effect to decrease or abolish dimer formation. For both Kirrel2 and Kirrel3, mutation of the same site, Q101 for Kirrel2 and Q128 for Kirrel3, to alanine abolished dimerization (Figure 4A and 4B). Alanine mutations of a set of equivalent positions at the dimerization interface of both Kirrels strongly diminished homodimerization: L58/L85, W69/Y96, R108/R135 (Kirrel2/Kirrel3 numbering). The energetic hot spot formed by these four residues create a continuous volume (Figure 4C) Surprisingly, the residue at the dimerization two-fold axis, Q52 for Kirrel2 and L79 for Kirrel3, which is likely important for specificity, is less crucial for binding energetics. In fact, the variable residues necessary for establishing dimerization specificity appear to cause smaller losses of binding energy when mutated, compared to the conserved residues at the hot spot. Overall, these mutagenesis experiments confirm that the interfaces observed in the crystal structures mediate dimer formation in solution.

**Figure 4.**
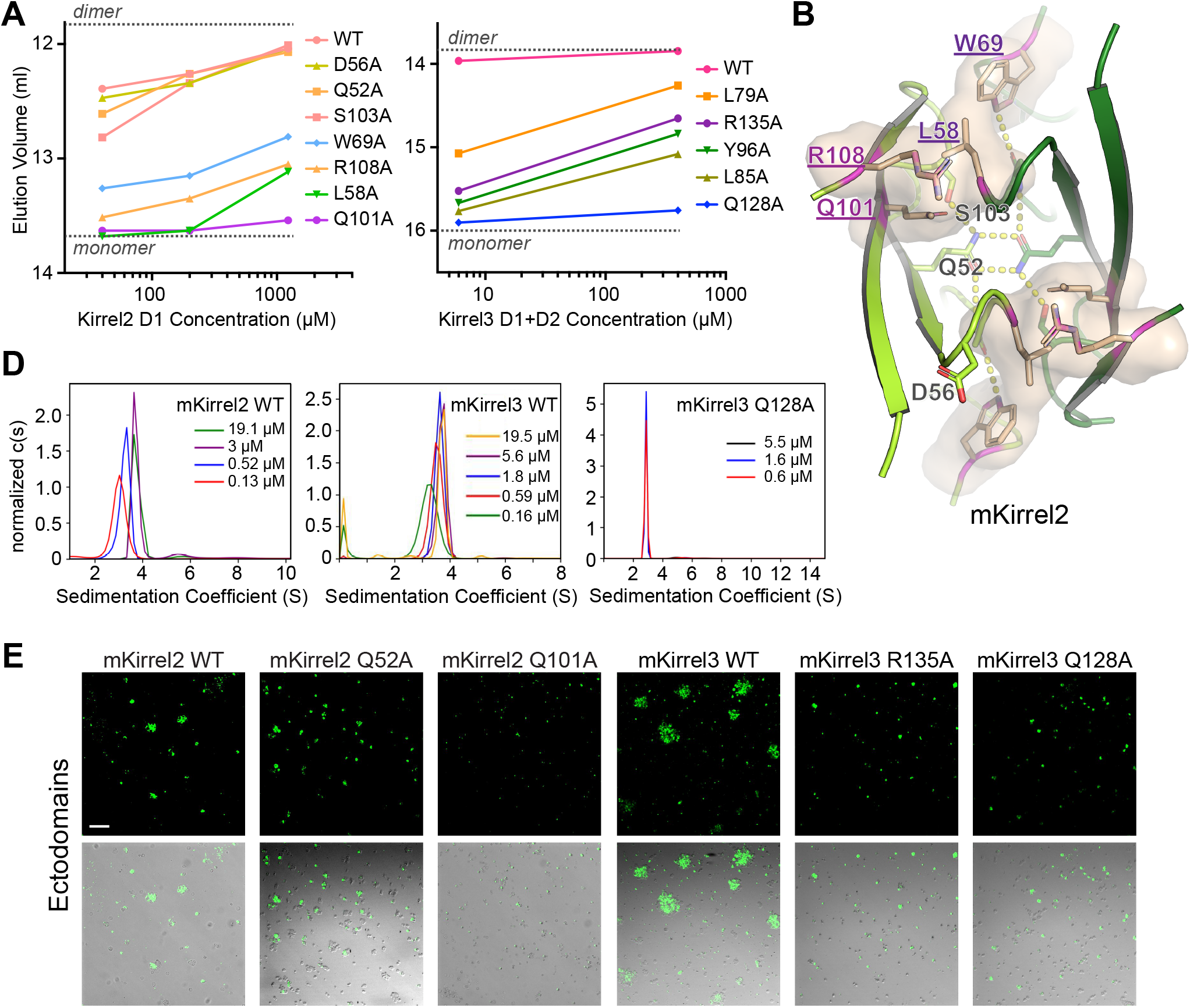
Mutational analysis of the Kirrel2 and Kirrel3 homodimerization interfaces. **A, B**. SEC elution volumes for wild-type and mutant Kirrel2 D1 (A) and Kirrel3 D1+D2 (B) loaded at multiple concentrations on the columns. A Superdex 75 10/300 column was used for the Kirrel2 D1 runs (left) and a Superdex 200 Increase 10/300 column was used for the Kirrel3 D1+D2 runs, both with a column volume of 24 ml. Expected monomeric and dimeric elution volumes are marked by dashed lines on the plots. See Figure S4 for SEC chromatograms. **C**. Four residues observed to be energetically important for Kirrel dimerization mapped onto Kirrel2 structure and shown as a surface (residue names purple and underlined). **D**. Sedimentation velocity results for mouse Kirrel2 WT (left), Kirrel3 WT (middle) and Kirrel3 Q128A (right) ectodomains performed at several initial protein concentrations, showing lack of dimerization for the Q128A mutant. KD for WT mKirrel2 and mKirrel3 refine to 0.9 µM [0.2 µM, 3.5 µM] and 0.21 µM [0.11 µM, 0.36 µM] (68.3% confidence intervals are shown in brackets). See Figure S4G,H for isotherms and fitting for dissociation constants. **E**. Cell aggregation assay for Kirrel2 and Kirrel3, WT and mutants, fused to intracellular GFP, performed with S2 cells. Scale bar, 100 µm.

### Kirrel2 and Kirrel3 homophilic adhesion is dependent on D1

While we have shown that the major dimerization interface for Kirrel homologs are within the N-terminal domain (D1) (Özkan et al., 2014, and above), it is not known if the other four domains can also cause dimerization in vertebrate Kirrels. In support of additional dimerization sites, Gerke et al. (2003) reported evidence that dimerization was not limited to any single domain within Kirrel1 and the related Nephrin. Furthermore, a large number of human mutations affecting Kirrel3 function in the ectodomain, but outside D1, have been identified (summarized in Taylor et al., 2020). To test whether Kirrel ectodomains can form dimers using interfaces we did not identify in our crystal structures, we used analytical ultracentrifugation to quantify dimerization of Kirrel2 and Kirrel3 ectodomains and one D1 interface mutant, Kirrel3 Q128A. We performed sedimentation velocity experiments at several concentrations and observed that both WT Kirrel ectodomains exist in a monomer-to-dimer equilibrium, confirming our SEC results (Figures 4D and S4G,H). Kirrel3 ectodomain with the D1 mutation Q128A showed no signs of dimerization at any concentration (Figure 4D), implying that no additional interface mediates dimerization.

Kirrel2 and Kirrel3 have been shown to mediate homophilic cell adhesion, which was suggested as necessary for their function in the regulation of olfactory sensory neuron axonal coalescence into glomeruli and synaptic specification in the hippocampus. We therefore examined the effect of non-dimerizing mutations on Kirrel-dependent adhesion using a cell aggregation assay. Suspension S2 cells were transfected with transmembrane constructs containing Kirrel ectodomains, attached to the transmembrane helix of Neurexin, followed by a cytoplasmic EGFP. We observed strong aggregation in transfected cultures, indicating that Kirrel2 and Kirrel3 form trans-homodimers that aggregate S2 cells (Figures 4E). When the Kirrel2 and Kirrel3 constructs carried mutations that abolish or strongly diminish dimerization through the D1 interface, such as Q128A in Kirrel3, cell aggregation was also abolished. However, an alanine mutation at the “specificity site” Kirrel2 Q52 did not abolish cell aggregation, as the soluble D1+D2 construct with this mutation still behaved as a dimer in SEC (Figure 4A). The analytical ultracentrifugation and cell aggregation experiments demonstrate that the D1 interface we observed in our crystal structures is required for Kirrel homodimerization, and a second interaction site within the ectodomains is either very weak to detect or does not exist.

### Kirrel ectodomains form elongated homodimers mediated by D1 interactions

As Kirrels are adhesion molecules carrying signaling motifs, the tertiary structure and conformational states of their ectodomains are relevant for their signaling and adhesive properties. To gain insights into the three-dimensional structure of Kirrel ectodomains, we studied Kirrel2 and Kirrel3 ectodomains in solution using an SEC-SAXS-MALS setup, including the Kirrel3 Q128A mutant (Figures 5 and S5, Table S2). For all ectodomains tested by SAXS (small-angle x-ray scattering), shapes of pair distance distributions, *P*(*r*), show strongly rod-like character, rather than globular or other shapes (Figures 5A, 5B, S5A and S5B). Kirrel2 and Kirrel3 homodimers have maximum dimensions (*D*_max_) of 39 and 35 nm, respectively, while Kirrel3 Q128A has a *D*_max_ of 21 nm, suggesting that Kirrel dimers are nearly double the length of the monomer. The accompanying MALS (multi-angle light scattering) data recapitulate our conclusions from pure SEC runs, where concentrated wild-type Kirrels have molar masses matching dimeric sizes and diluted Kirrel3 samples yield intermediate values between predicted monomer and dimer masses, while Kirrel3 Q128A is a pure monomer (Figure S5C and Table S3). These results agree with our elongated Kirrel model where only D1 domains interact between Kirrel monomers, and support a D1-D1, or N-terminal tip-to-tip, interaction model of Kirrel homodimerization.

**Figure 5.**
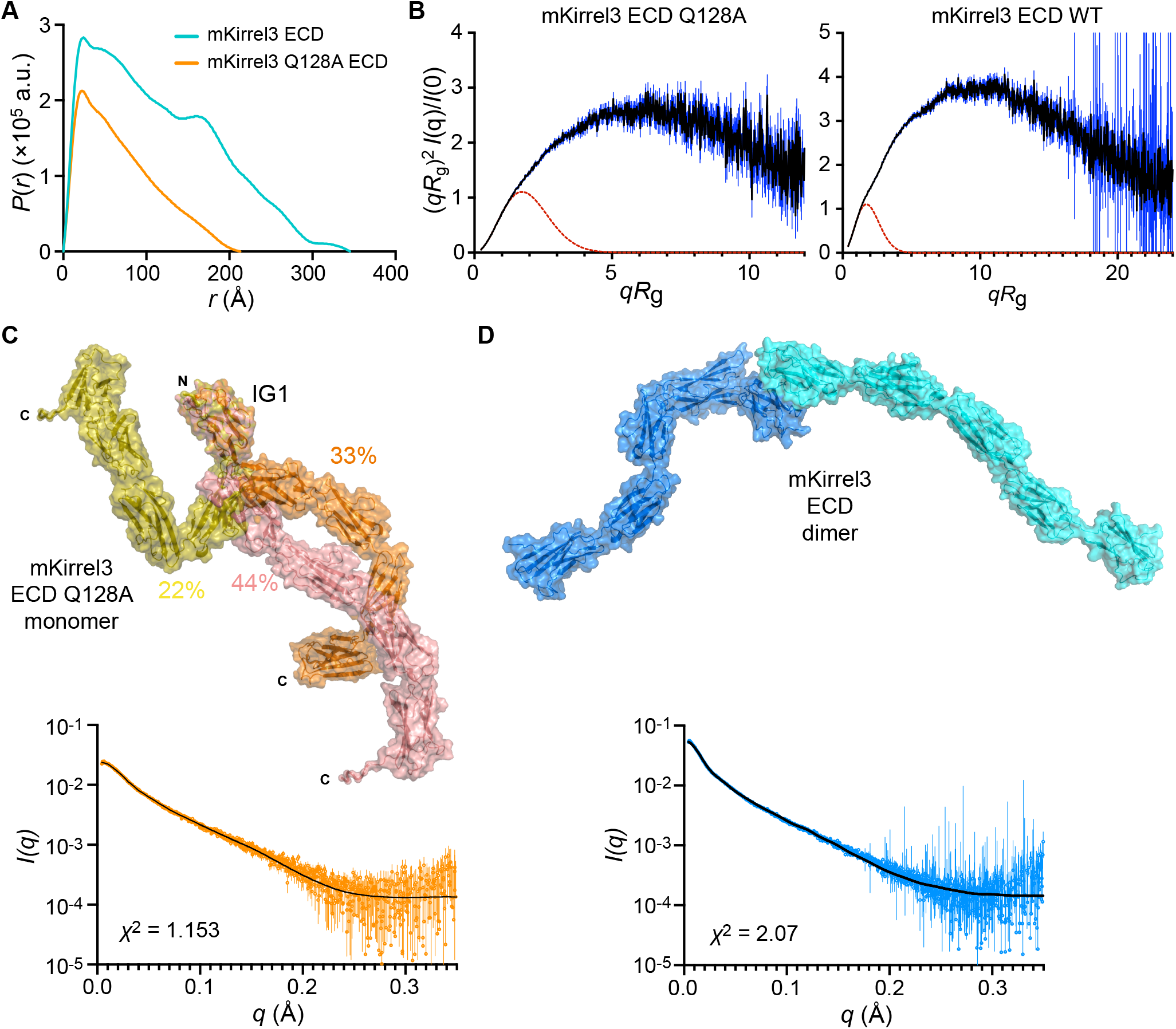
SAXS analysis of Kirrel ectodomains. **A**. Pair distance distributions, *P*(*r*), for mKirrel3 ectodomains, WT and Q128A. WT Kirrel3 is a longer molecule than Kirrel3 Q128A since it has a larger *D*_max_ (35 nm vs. 21 nm). **B**. Dimensionless Kratky plots for mKirrel3 ectodomains; blue vertical lines are measured errors. Dashed red lines show predicted plots for rigid, globular molecules with the same *R*_g_. **C**. Ensemble model fitting of SAXS data for mKirrel3 Q128A ectodomain using *EOM*. The three-model ensemble with contributions are indicated as percentages next to the models. The scattering profile predicted from the ensemble model (black) closely matches observed scattering data (measurement errors are indicated as orange vertical bars). **D**. *SASREF* model of mKirrel3 WT ectodomain dimer, and its predicted scattering (black) overlayed on observed SAXS data (cyan) with vertical bars representing measurement errors.

A prominent feature common to Kirrel2, Kirrel3, and Kirrel3 Q128A ectodomains observed in dimensionless Kratky plots for SAXS data (Figures 5B and S5B) is flexibility, likely reflecting significant interdomain movements. While this may complicate low-resolution modeling of the ectodomains, we used *DAMMIF* (Franke and Svergun, 2009) to create ab-initio bead models. Bead models for Kirrel2 and Kirrel3 appear as a thin V with a vertex angle of approximately 115°, while a bead model for Kirrel3 Q128A is a straight rod that is as long as each of the “wings” of the Kirrel2 and Kirrel3 models (Figures S5D and E). These conformations agree with our structural models of a fixed angle of Kirrel2 and Kirrel3 dimerization defined by the D1-D1 binding interface.

Next, we attempted to model Kirrel ectodomains using an ensemble representation with *EOM* from the *ATSAS* suite (Tria et al., 2015), where flexibility was allowed between five rigid-body-defined IG domains, which were available through our high-resolution Kirrel D1 structures or were modeled (Domains 2 to 5) based on similar immunoglobulin domain templates. *EOM* can explain observed SAXS profiles for the monomeric Kirrel3 Q128A using a three-model ensemble that has theoretical scattering matching observed data with high fidelity (*χ*^2^ = 1.15, Figure 5C). While the homodimer also shows strong flexibility based on the Kratky analysis (Figure 5B), ensemble modeling did not yield an equally satisfactory model for explaining WT Kirrel3 scattering data, possibly due to the presence of monomeric species in solution and a higher-degree of flexibility caused by movements between ten, rather than five domains. For the dimeric Kirrel3 WT SAXS data, we used *SASREF* from the same package to allow for interdomain flexibility within one model (Petoukhov and Svergun, 2005) (Figure 5D). When the resulting model was aligned to the SAXS bead model from *DAMMIF*, we observed a strong overlap, supporting the validity of our SAXS bead models as well as our model of Kirrel tip-to-tip dimerization (Figures S5D and E). These results further point to a lack of secondary dimerization sites.

### Kirrel3 Q128A mutation causes defects in glomerulus formation in the AOB

In order to assess whether the Kirrel D1-to-D1 model of homophilic adhesion contributes to Kirrel3 function in wiring the nervous system, we examined the effect of abolishing Kirrel3 dimerization on vomeronasal sensory neuron axon targeting in the AOB. We engineered mice expressing the Kirrel3 Q128A amino acid substitution, which abolishes Kirrel3 dimerization, using CRISPR technology (Figure 6A-C). An analysis of Kirrel3 protein in lysates extracted from brain samples of wild-type and *Kirrel3*^*Q128A/Q128A*^ mice reveals that Kirrel3 Q128A is expressed at comparable levels to the wild-type protein in the brain (Figure 6D,E). Additionally, cell surface biotinylation assays on brain slices from control and *Kirrel3*^*Q128A/Q128A*^ mice, combined with flow cytometry of HEK293T cells transfected with full-length WT and Q128A Kirrel3, reveals that the mutant protein is properly trafficked to the cell’s surface (Figure 6F and Figure S6).

**Figure 6.**
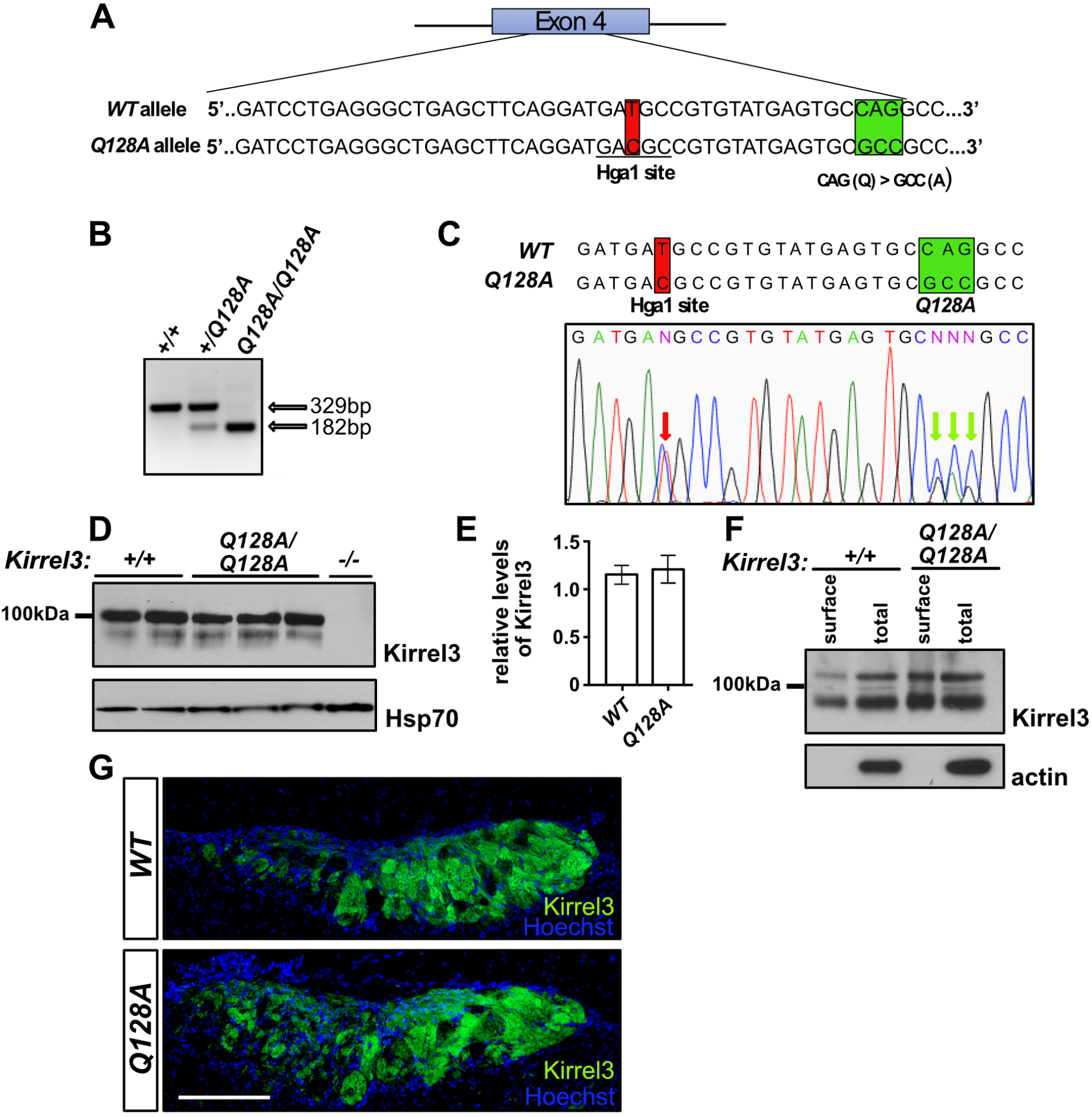
Characterization of the Kirrel3 Q128A mouse. **A**. Diagram of the generation of the *Kirrel3 Q128A* mouse. Mice carrying a modified *Kirrel3* allele containing mutations that modify amino acid 128 from a glutamine (Q) to an alanine (A), as well as a silent mutation introducing an HgaI restriction enzyme cutting site for genotyping purposes, were generated. Green square: Q to A mutations. Red square: mutation creating an HgaI restriction site. **B-C**. Identification of the *Kirrel3 Q128A* allele by restriction enzyme digest and DNA sequencing. Digestion with HgaI (B) and DNA sequencing (C) of a PCR fragment from exon 4 of the *Kirrel3* allele demonstrate the presence of the newly introduced HgaI restriction site. DNA sequencing also reveals the presence of the three nucleotide substitutions resulting in the Q to A amino acid substitution in a *Kirrel3*^*+/Q128A*^ mouse. The red arrows in the electropherogram in C indicate the overlapping peaks caused by the nucleotide substitutions in one of the Kirrel3 alleles. **D**. Quantification of Kirrel3 protein by western blot of brain lysate collected from *Kirrel3*^*+/+*^ and *Kirrel3*^*Q128A/Q128A*^ mice shows that similar levels of Kirrel3 and Kirrel3 Q128A levels are expressed in the brain of these mice, respectively. Data were analyzed using unpaired *t*-test; *n*=3 for *Kirrel3*^*+/+*^ and 4 for *Kirrel3*^*Q128A/Q128A*^ mice. **E**. Surface membrane distribution of Kirrel3 Q128A in acute brain slices. Western blots of acute brain slice lysate collected from *Kirrel3*^*+/+*^ and *Kirrel3*^*Q128A/Q128A*^ mice following incubation with biotin and isolation of surface proteins by batch streptavidin chromatography. Both Kirrel3 and Kirrel3 Q128A are distributed to the cell surface. **F**. Immunohistochemistry on sagittal sections of the AOB from *Kirrel3*^*+/+*^ and *Kirrel3*^*Q128A/Q128A*^ adult mice labelled with antibodies against Kirrel3 and Hoechst. The Kirrel3 and Kirrel3 Q128 proteins can be detected in subsets of glomeruli in both the anterior and posterior regions of the AOB in *Kirrel3*^*+/+*^ and *Kirrel3*^*Q128A/Q128A*^ mice, respectively. Scale bar is 200 μm.

We then investigated the expression patterns of Kirrel3 Q128A on VSN axons innervating the AOB. Kirrel3 is expressed on subsets of VSN axons innervating glomeruli throughout the AOB with a large proportion of glomeruli in the posterior region expressing Kirrel3 (Prince et al., 2013). Sagittal sections of the AOB stained for Kirrel3 demonstrated a clear localization of Kirrel3 in a majority of glomeruli in the posterior region of the AOB in both wild-type and *Kirrel3*^*Q128A/Q128A*^ mice (Figure 6G). Since *Kirrel3*^*-/-*^ mice show altered glomeruli structure in the posterior region of the AOB (Prince et al., 2013), we next performed a detailed analysis of the glomerular layer in *Kirrel3*^*Q128A/Q128A*^ mice. To visualize and delineate glomeruli, sagittal sections of the AOB of control and *Kirrel3*^*Q128A/Q128A*^ mice were stained with anti-VGLUT2, which marks excitatory pre-synaptic terminals in the AOB. As previously observed in *Kirrel3*^*-/-*^ mice (Prince et al., 2013), the size and number of glomeruli in the anterior region of the AOB appears unaffected in *Kirrel3*^*Q128A/Q128A*^ mice when compared to control animals (Figures 7 A, B, G, I). In contrast, and as observed in *Kirrel3*^*-/-*^ mice, the posterior region of the AOB in *Kirrel3*^*Q128A/Q128A*^ mice contains significantly fewer glomeruli and the average size of each glomerulus in increased by ∼43% (Figures 7 A-C, H, J). Thus, the Kirrel3 D1-D1 binding interface is necessary for proper coalescence of VSN axons into glomeruli of the AOB.

**Figure 7.**
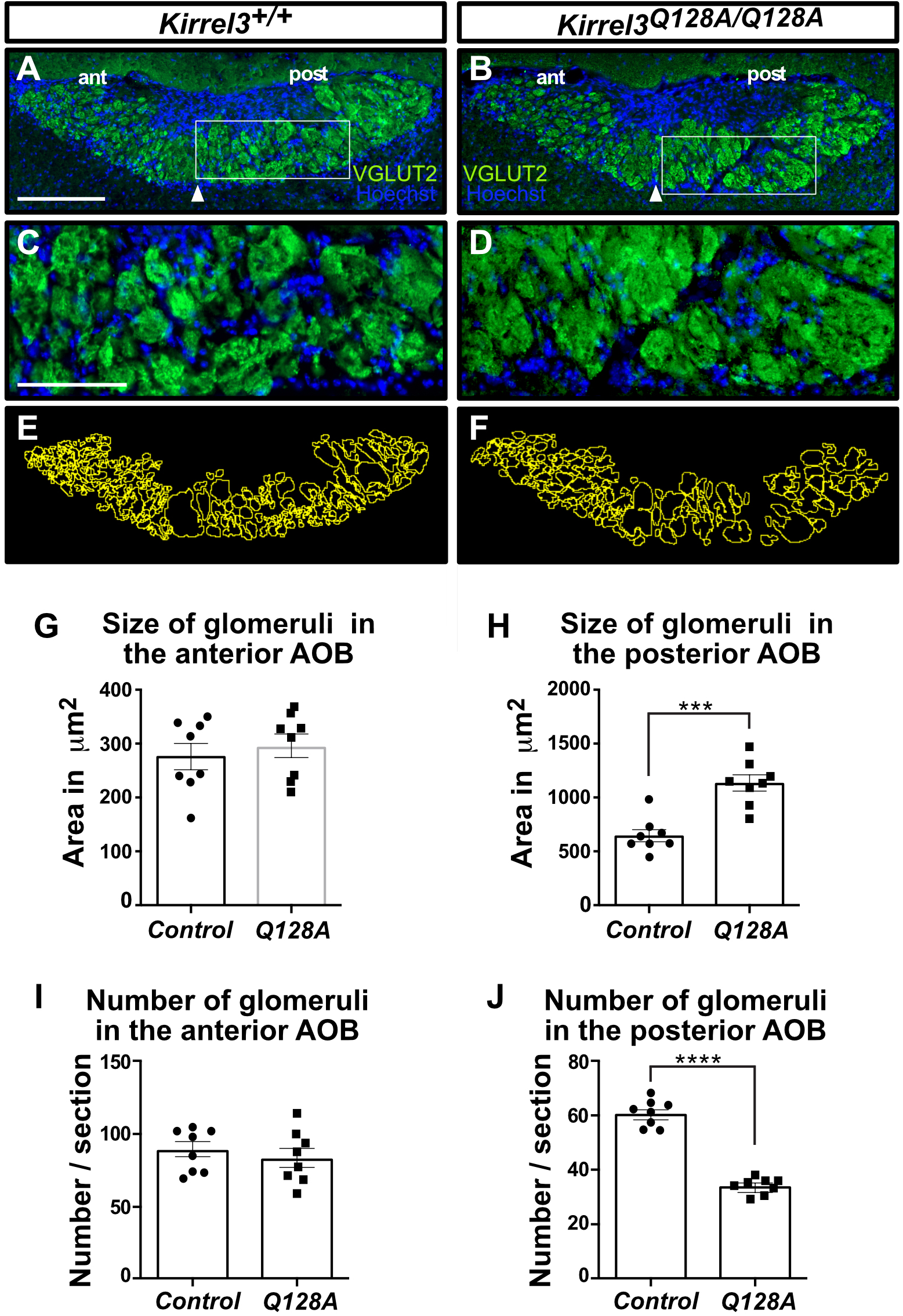
Glomeruli structure is altered in the AOB of *Kirrel3*^*Q128A/Q128A*^ mice. **A-F**. Parasagittal sections of the AOB from adult *Kirrel3*^*+/+*^ (A,C) and *Kirrel3*^*Q128A/Q128A*^ (B,D) mice, labelled with a VGLUT2 antibody and Hoechst (A-D). Higher magnification of outlined regions in A and B are shown in C and D, respectively. Glomeruli in the posterior region of the AOB in *Kirrel3*^*Q128A/Q128A*^ mice appear significantly larger and less numerous when compared to *Kirrel3*^*+/+*^ mice (E,F). Arrowheads denote the boundary between the anterior (ant) and posterior (post) regions of the AOB. **E-J**. Quantification of the size and number of glomeruli in the anterior (G,I) and posterior regions (H,J) of the AOB in adult control and *Kirrel3*^*Q128A/Q128A*^ mice. A representation of the glomeruli outlining approach used for quantification using sections in panels A and B as examples are shown in E and F, respectively. A significant increase in the size of glomeruli in the posterior (*Control*: 644.0 ± 56.2 µm^2^; *Kirrel3*^*Q128A/Q128A*^: 1134.1 ± 74.5 µm^2^), but not anterior (*Control*: 284.0 ± 19.6 µm^2^; *Kirrel3*^*Q128A/Q128A*^: 295.6 ± 22.1 µm^2^), region of the AOB is observed in the *Kirrel3*^*Q128A/Q128A*^ mice (G,H). There is also a decrease in glomeruli numbers in the posterior (*Control*: 61.15 ± 1.75; *Kirrel3*^*Q128A/Q128A*^: 34.27 ± 1.02), but not anterior (*Control*: 87.21 ± 4.92; *Kirrel3*^*Q128A/Q128A*^: 84.22 ± 6.33), region of the AOB in *Kirrel3*^*Q128A/Q128A*^ mice (J). Data were analyzed using unpaired t-test; *n*=8 mice for each genotype. \*\*\*\**p*-value < 0.0001 (glomerular counts), \*\*\**p*-value < 0.001 (glomerular size), error bars: ± SEM. Scale bars: 250 µm in A,B and 100 µm in C,D.

## DISCUSSION

The wiring of the nervous system is guided by a combination of intercellular interactions mediated by cell surface receptors (Sanes and Zipursky, 2020). The interactions can be heterotypic, usually allowing for establishing connectivity between different neuronal types, guiding growth of axons in a gradient field established by cues or by contacts between processes resulting in attraction or repulsion. Homotypic interactions can also be used to establish synaptic connections (as Kirrel3 is known to mediate in the hippocampus), lead to contact-mediated repulsion or to fasciculation and coalesce axons. Kirrel2 and Kirrel3 are differentially expressed in sets of sensory neurons whose axons are segregated into separate glomeruli in the accessory and main olfactory bulbs. The molecular mechanism through which differential expression of Kirrel family members in populations of sensory axons modulates their segregation into glomeruli remained to be addressed.

Structures presented here show that Kirrel2 have key residues that allow the formation of a hydrogen bonding network at the homodimeric interface, while Kirrel3 evolved a hydrophobic interface in lieu of this hydrogen bonding network. Ancestral vertebrate Kirrel has crucial amino acids compatible with the presence of the hydrogen bonding network, and this specialization of a non-polar Kirrel3 interface core likely evolved to ensure exclusive homodimerization of the two Kirrels, and no heterodimerization, allowing for the segregation of OSN termini into separate Kirrel2 and Kirrel3-expressing glomeruli. This finding of incompatible interface chemistries is distinct to the vertebrate Kirrels: The two fruit fly Kirrel dimerization interfaces show strong conservation and compatible chemistries that would allow them to create putative complexes (Özkan et al., 2014), and there is evidence that fly Kirrels can form heterodimers (Özkan et al., 2013). We otherwise found that Kirrel duplications outside vertebrates is relatively rare.

The distribution of extant Kirrel proteins we identified in cyclostomes and gnathostomes are consistent with the duplication of Kirrels during the early chordate whole-genome duplication events (Dehal and Boore, 2005). Given that ancestral Kirrel2 and Kirrel3 have chemically incompatible interface sequences similar to mouse Kirrels and that extant jawed fish Kirrel sequences also follow these patterns, Kirrel specialization must have appeared early in gnathostomes and before the rise of distinct accessory olfactory systems and several expansions in olfactory receptor gene families (Bear et al., 2016; Poncelet and Shimeld, 2020). Therefore, it is attractive to speculate that specialization of Kirrels and other cell surface receptors differentially expressed in OSNs and VSNs and duplicated during the whole-genome duplication events, may have set the stage for increases in ORs, OSNs and glomeruli in main and accessory olfactory bulbs.

Kirrels have been identified and implicated in numerous functional contexts in the nervous system. Loss of Kirrel2 and Kirrel3 causes disorganization of a subset of glomeruli in the main and accessory olfactory bulbs (Brignall et al., 2018; Nakashima et al., 2019; Prince et al., 2013; Vaddadi et al., 2019), and Kirrel3 knockout mice show behavioral abnormalities similar to those in autistic patients, including defects in social responses, auditory sensory processing and communication, and repetitive behaviors (Hisaoka et al., 2018; Völker et al., 2018), in addition to loss of male-male aggression likely as a result of miswiring in the AOB (Prince et al., 2013). Furthermore, several mutations in human autism and intellectual disability patients were identified (see Taylor et al., 2020 for a list). To our surprise, none of the missense mutations are in the first domain, including mutations recently reported to affect Kirrel3-mediated cell adhesion (Taylor et al., 2020). We therefore studied the entire Kirrel2 and Kirrel3 ectodomains using a combination of biophysical methods, but could not produce evidence that there are secondary binding sites outside the D1-D1 interface. A single alanine mutation at the D1-D1 Kirrel3 dimer interface completely abolishes ectodomain dimerization at micromolar concentrations. SAXS data suggest highly elongated models for dimerized Kirrels compatible with D1-D1 intermolecular interactions, further rendering other putative binding sites unlikely or very weak. Additional structural studies on liposomes and lipid bilayers may be necessary to refute or confirm secondary *cis* or *trans* binding sites.

We also showed that the dimerization interface observed in our structures directly contributes to glomerulus formation, at least in the case of Kirrel3. Morphological analysis of glomeruli in the AOB using the Kirrel3 Q128A substitution closely resembled those of the *Kirrel3* knockout, where glomeruli increase in size but decrease in number in the posterior AOB. We have confirmed that the Q128A mutant displays correctly to the cell surface, which implies that the observed AOB phenotype is not due to misfolding or trafficking errors. Though we did not test Kirrel2 loss-of-function mutations *in vivo*, Kirrel2 shares a closely related structure and evolutionary history with Kirrel3, and it is reasonable to assume that the dimerization interface on Kirrel2 D1 may also control glomerulus formation in a similar manner. However, while Kirrel homodimerization is necessary for proper glomerulus formation, it is likely that other molecules contribute to glomerular segregation of axons, including expression of vomeronasal receptors and other receptor pairs (such as Eph-Ephrin), and neuronal activity.

Multiple lines of evidence support the idea that Kirrels and its homologs act as more than “molecular Velcro” in nervous system development. In the hippocampus, Kirrel3 homophilic adhesion is required, but not sufficient for synapse formation, and mutations that occur outside of the D1 binding domains and even in the intracellular domains prevent synapse formation, despite being able to form homodimeric complexes (Taylor et al., 2020). The *C. elegans* homologs SYG-1 and SYG-2 were shown to be signaling receptors, whose functions depend on the exact geometry of ectodomain dimerization also mediated by D1 (Özkan et al., 2014). Finally, Kirrels carry conserved and functional signaling motifs in their cytoplasmic domain, including phosphorylation sites, PDZ motifs, and actin cytoskeleton recruitment sequences (Bulchand et al., 2010; Chia et al., 2014; Gerke et al., 2006; Harita et al., 2008; Huber et al., 2003; Sellin et al., 2003; Yesildag et al., 2015). These findings suggest that Kirrel3 undergoes active signaling once the homodimeric complexes have formed, which may be influenced by the overall flexibility and conformation of the extracellular domains and how tightly Kirrels may pack at the site of a Kirrel-mediated cell adhesion.

The question remains, however, of whether Kirrel2 and Kirrel3 also actively signal for the formation of glomeruli, and may also regulate synapse formation of sensory neurons in these glomeruli with second-order neurons, such as mitral cells. As Kirrel3 has been shown to specify synapse formation in the hippocampus between dentate gyrus and CA1 neurons, it is possible that Kirrels play a dual role of simple adhesion and synapse specification. Some second-order neurons in the AOB were observed to selectively form dendrites to VSNs expressing the same vomeronasal receptors (Del Punta et al., 2002), indicating that synapses formed in the glomeruli may be specified by cell surface receptors controlled by vomeronasal receptor expression, such as Kirrels, to organize sensory input. A mechanism through which Kirrels mediate both VSN coalescence and synapse formation onto mitral cells would be appealing for its simplicity in explaining wiring specificity.

## ACKNOWLEDGMENTS

We thank Nathan Canniff for technical support, Davide Comoletti for suggestions and guidance, and the McGill Integrated Core for Animal Modeling supported by the Goodman Cancer Research Centre for their advice and generation of genetically-modified animals. We acknowledge excellent support by Chad Brautigam with analytical ultracentrifugation experiments at the UT Southwestern Macromolecular Biophysics Resource Laboratory, and Srinivas Chakravarthy at the Advanced Photon Source (APS) SAXS beamline BioCAT 18-ID. This research used resources of the APS, a U.S. Department of Energy (DOE) Office of Science User Facility operated for the DOE Office of Science by Argonne National Laboratory under Contract No. DE-AC02-06CH11357. Work at BioCAT was supported by grant 9P41GM103622 from the National Institute of General Medical Sciences (NIGMS) of the National Institutes of Health (NIH). The Pilatus 3 1M detector in BioCAT was provided by grant 1S10OD018090-01 from NIGMS. X-ray diffraction data were collected at the Stanford Synchrotron Radiation Lightsource (SSRL) beamline 12-2 and APS beamline 24-ID-E. Use of the SSRL, SLAC National Accelerator Laboratory, is supported by the U.S. DOE, Office of Science, Office of Basic Energy Sciences under Contract No. DE-AC02-76SF00515. The SSRL Structural Molecular Biology Program is supported by the DOE Office of Biological and Environmental Research, and by the NIH, NIGMS (including P41GM103393). Beamline 24-ID-E at the Northeastern Collaborative Access Team is funded by NIGMS, NIH (P30 GM124165). We acknowledge funding support from NIH through grants R01NS097161 (to E.Ö.), from the Canadian Institutes of Health Research and the Natural Sciences and Engineering Council of Canada (to J.F.C.), from the Healthy Brain Healthy Lives program of McGill University and Fonds de Recherche du Québec – Santé (to N.V. and S.Q.). Y.P. was supported by Samsung Scholarship and J.S.P. was supported by a Steiner Award from the University of Chicago’s Division of Biological Sciences.

## AUTHOR CONTRIBUTIONS

E.Ö. and J.F.C. conceptualized the study. J.W. and J.S.P. designed and performed the cell biology and biophysical experiments. C.A.R. and E.Ö. determined crystal structures. Y.P., E.Ö. and J.W.T. performed the phylogenetic analysis. N.V., S.Q., and E.D. were involved in the generation and characterization of transgenic animals. J.S.P., N.V., E.Ö. and J.F.C. wrote the manuscript.

## METHODS

### Protein Expression and Purification

For biophysical and structural studies, mouse Kirrel2 and Kirrel3 constructs were expressed in High Five cells using the baculovirus expression system. All expression constructs were tagged C-terminally with hexahistidine tags in the baculoviral transfer vector pAcGP67A (BD Biosciences). Proteins were purified using Ni-NTA agarose (Qiagen) resin first, followed by size-exclusion chromatography with Superdex 75 10/300 or Superdex 200 10/300 Increase columns (GE Healthcare) in 1x HBS (10 mM HEPES pH 7.2, 150 mM NaCl). We observed that some of the Kirrel constructs tend to precipitate at higher concentrations at 4°C; therefore, proteins were purified and stored only for short periods at 16 to 22°C.

### Protein Crystallization

Purified mKirrel2 D1 was concentrated to 10.6 mg/ml and crystallized using the sitting-drop vapor-diffusion method in 0.4 M Potassium/Sodium tartrate. Crystals were cryo-protected in 0.5 M Sodium acetate, 30% Ethylene glycol, and vitrified in liquid nitrogen. Complete diffraction dataset was collected at the SSRL beamline 12-2.

Purified mKirrel3 D1 was concentrated to 9.1 mg/ml and crystallized using the sitting-drop vapor-diffusion method in 0.1 M Sodium cacodylate, pH 6.5, 1.4 M Sodium acetate. Crystals were cryo-protected in 0.1 M Sodium cacodylate, pH 6.5, 1.4 M Sodium acetate with 30% Ethylene glycol, and vitrified in liquid nitrogen. Complete diffraction dataset was collected at the APS beamline 24-ID-E.

### Structure Determination by X-ray Crystallography

Diffraction data were reduced using *HKL2000* (Otwinowski and Minor, 1997). Structure of mKirrel2 D1 was phased with molecular replacement using mouse Duf/Kirre D1 as model (PDB ID: 4OFI; Özkan et al., 2014) with *PHASER* (McCoy et al., 2007) as part of the *PHENIX* suite (Liebschner et al., 2019). For molecular replacement of the mKirrel3 D1 data set, a partly refined mKirrel2 D1 structure was used. Model building and refinement were done using *Coot* (Emsley et al., 2010) and *phenix*.*refine* (Afonine et al., 2012), respectively. mKirrel3 D1 model was refined with eight TLS groups, one for each chain in the asymmetric unit, while no TLS refinement was used for the mKirrel2 model. Crystal structures and structure factors are deposited with the Protein Data Bank with accession codes 7LTW (mKirrel2 D1) and 7LU6 (mKirrel3 D1). Crystallographic data and refinement statistics are tabulated in Table S1.

### Phylogenetics and Ancestral Sequence Reconstructions

Putative Kirrel orthologs were identified using a reciprocal *BLAST* strategy with the three mouse Kirrels, mouse Nephrin, and *D. melanogaster* Rst and Kirre. Each sequence hit was confirmed to contain conserved features of Kirrels and Nephrins, specifically five and ten IG/FnIIII domains, respectively, in the extracellular regions, and a membrane helix. A total of 90 Kirrel and 41 Nephrin-like proteins were identified across major protostome and deuterostome taxa; we failed to detect clear Kirrel and Nephrin orthologs in other metazoan groups.

The mature ectodomains of 90 Kirrel sequences were aligned using *MUSCLE* (version 3.8.425) with default parameters (Edgar, 2004), as the other regions cannot be aligned with high confidence. Parts of the alignment with insertions represented by few sequences were removed, and maximum likelihood phylogenies were inferred using *RAxML-NG* v1.0.1 (Kozlov et al., 2019) using the best-fit LG+I+G4 evolutionary model determined by *MODELTEST-NG* (Darriba et al., 2020). Among the three vertebrate Kirrels, Kirrel2 shows faster and heterogenous rates of evolution in gnathostomes. Ancestral sequences were inferred using the marginal reconstruction algorithm in *PAML* v4.8 (Yang, 2007) using the LG+G4 model and the amino acid frequencies inferred on the ML tree by *RAxML* v8.2.12.

### Analytical Ultracentrifugation

Analytical ultracentrifugation data was collected at the UT Southwestern Macromolecular Biophysics Resource laboratory. mKirrel2 and mKirrel3 ectodomain samples were placed in 0.3 or 1.2-cm charcoal-filled Epon centerpieces sandwiched between sapphire windows. Cells were placed in an An50-Ti rotor and spun at 20°C at 50,000 rpm for sedimentation velocity experiments. During the spins, concentration profiles were measured using absorbance at 230 nm or using Rayleigh interferometry for the highest concentration samples (blue points in isotherms in Figures S4G,H). The data were analyzed using the *c*(*s*) methodology in *SEDFIT* (Schuck, 2000). Partial-specific volume, density, and viscosity were calculated using *SEDNTERP* (Laue et al., 1992), and the following values were used in data analysis: *ε*_230_ = 389,836 (mKirrel2) and 395,466 (mKirrel3) M^-1^cm^-1^; *ε*_interferometric_ = 150,896 (mKirrel2) and 151,329 (mKirrel3) M^-1^cm^-1^; *ρ*(solution) = 1.0052 g/cm3; *η*(solution) = 0.010249 Poise.

To calculate dissociation constants, *GUSSI* (Brautigam, 2015) was used to integrate the *c*(*s*) distributions, which were assembled into isotherm files that were imported into *SEDPHAT*, where a monomer-dimer model was imposed (Schuck, 2003) with a fixed *s*-value for the monomer (2.87 S). All AUC figures were rendered in *GUSSI*.

The Q128A mutant for the mKirrel3 ectodomain displayed an unmoving peak at all concentrations with a calculated sedimentation coefficient of 2.87 S, which represents the monomer, while the WT mKirrel2 and mKirrel3 ectodomains had c(s) distributions showing increasing sedimentation coefficients with increasing protein concentrations, where dimeric mKIrrel2 and mKirrel3 have refined sedimentation coefficients of 3.9 and 3.8 S, respectively. The single peaks observed in the *S* vs. *c*(*s*) plots, in addition to size-exclusion chromatography elution profiles showing single peaks corresponding to intermediate hydrodynamic sizes, strongly suggest fast rates of dimerization kinetics (*k*_off_> 10^−3^ s^-1^)

### Cell Aggregation Assays

Kirrel ectodomains were cloned in the Drosophila expression plasmid pAWG (Drosophila Genomics Resource Center), and transiently transfected in S2 cells using Effectene (QIAGEN, catalog no. 301425). Cells were collected three days post-transfection, spun down at 250 x g for 5 minutes, washed once with phosphate-buffered saline, and re-suspended in serum-free Schneider’s media (Lonza, catalog no. 04-351Q) at 10^7^/ml cell density before imaging.

### SEC-SAXS-MALS

Scattering data was collected for Kirrel ectodomains at the Advanced Photon Source beamline 18-ID using a SEC-SAXS-MALS setup. *BioXTAS RAW* version 2.0.2 was used for SAXS data collection and data reduction (Hopkins et al., 2017). The *RAW* interface was also used for Guinier analysis of the scattering data and the indirect Fourier transform methods implemented in *GNOM* (Svergun, 1992), part of the *ATSAS* package (Manalastas-Cantos et al., 2021), for pair-distance distribution analysis. Domains 2 to 5 for mKirrel3 were modeled using the I-TASSER server (Roy et al., 2010; Yang and Zhang, 2015). Modeling using the SAXS data was performed using the *ATSAS* package version 3.0.3 with the *ab initio* shape determination by bead modeling in *DAMMIF* (*Franke and Svergun, 2009*), with the ensemble optimization method implemented in *EOM* (Tria et al., 2015), and with the simulated annealing protocol for connected domains in *SASREF* (Petoukhov and Svergun, 2005). Further details about SAXS data collection and analysis are tabulated in Table S2.

Light scattering data was collected at the APS 18-ID beamline on a DAWN HELEOS with a T-rEX refractometer (Wyatt Technology) at 25°C, and analyzed in *ASTRA* version 7.3.2.19. *dn*/*dc* values of 0.181 to 0.182 were used to account for predicted glycosylation for molar mass calculations. Further details about MALS data collection and analysis are tabulated in Table S3. Expected molecular masses for Kirrel2 and Kirrel3 WT ectodomain constructs without glycosylation are both 53.0 kDa. With predicted N-linked glycosylation added by the insect cell expression system, we expect molecular masses for Kirrel2 and Kirrel3 WT ectodomain monomers to be 56 to 59 kDa.

### Flow Cytometry Assay for Cell Surface Trafficking

Kirrel3 WT and Q128A cDNA, excluding the signal peptide, were cloned C-terminal to a preprotrypsin leader sequence and a FLAG peptide (DYKDDDDK) under the control of the CMV promoter, and transfected into HEK293T cells using LipoD293 (SignaGen Laboratories, catalog no. SL100668). After 48 h, transfected cells were detached from the cell culture plate by incubating cells in a citric saline solution (135 mM potassium chloride, 50 mM sodium citrate) at 37°C for 5 min, followed by addition of Dulbecco’s modified Eagle medium (Thermo Fisher, catalog no. 11965092) with 10% fetal bovine serum and mechanical suspension of cells. Cells were spun down at 500 *g* for 5 min washed with phosphate-buffered saline (PBS; Thermo Fisher, catalog no. BP661-50, 137 mM NaCl, 2.7 mM KCl, 11.9 mM phosphate, pH 7.4), then resuspended in PBS + 0.1% bovine serum albumin (BSA). For staining of Kirrel3 displayed on the cell surface, cells were incubated with anti-FLAG M2 antibody (Sigma, catalog. F3165; 1:1000 dilution in PBS with 0.1% BSA) for 30 min at room temperature, washed twice with PBS + 0.1% BSA, then incubated with a secondary donkey anti-mouse Alexa Fluor 488 antibody (Thermo Fisher, catalog no. A-21202; 1:500 dilution in PBS with 0.1% BSA). In preparation for flow cytometry, cells were washed twice and resuspended in PBS + 0.1% BSA. Surface display of the Kirrel3 ectodomains on HEK cells were measured on the Accuri C6 flow cytometer (BD Biosciences) with the 488 nm excitation laser and the 533 nm filter in 10,000 cells per sample.

### Generation of *Kirrel3 Q128A* mouse line

sgRNAs were designed using the open access software Breaking-Cas (http://bioinfogp.cnb.csic.es/tools/breakingcas) and tested for cutting efficiency using a T7 assay as previously described (Sakurai et al., 2014). One-cell stage C57Bl6 mouse zygotes were microinjected with the sgRNA, Cas9 protein, and a donor DNA template to introduce the required DNA mutations in the *Kirrel3* allele (see Figure 6A) and were implanted in surrogate female mice. Offsprings were screened for the presence of the target mutations by PCR, restriction enzyme analysis, and DNA sequencing. Three *Kirrel3 Q128A* founder lines were chosen for further analyses and backcrossed for three generations in the C57Bl6 background to segregate any potential off-target mutations. Analyses of the glomerular structure of the AOB presented in Figure 7 was performed on *Kirrel3 Q128A* mice derived from a single founder line. Similar analyses were also performed on *Kirrel3 Q128A* mice derived from two other founder lines and revealed similar phenotypes (data not shown). All animal procedures have been approved by The Neuro Animal Care Committee and McGill University, in accordance with guidelines of Canadian Council of Animal Care.

### Immunohistochemistry

Two to three-month-old adult mice were anesthetized and perfused transcardially with 10 ml ice-cold 1x PBS followed by 10 ml 4% paraformaldehyde solution in 1x PBS. Brains were dissected and post-fixed in 4% paraformaldehyde solution for 30 min followed by 24 h cryoprotection in 30% sucrose. 20 µm-thick sagittal AOB sections were collected on microscope slides and incubated overnight at 4°C with the following primary antibodies: Anti-VGLUT2, 1:500 (Synaptic Systems) or anti-Kirrel3, 1:100 (Neuromab). After rinsing in Tris-Buffered Saline, the appropriate secondary antibody-Alexa 488 conjugate (Molecular Laboratories) was applied at 1:500 dilution to detect the primary antibody and BS lectin at 1:1500 (Vector Laboratories) was applied along with the secondary antibody. Sections were counter-stained with Hoechst at 1:20,000 dilution (Molecular Probes).

### Analysis of Glomeruli in the AOB

Analysis of the size and number of glomeruli in AOB sections were performed as previously described (Brignall et al., 2018; Prince et al., 2013). 20 µm-thick sagittal sections of the AOB were obtained from *Kirrel3*^*+/+*^, *Kirrel3* ^*+/Q128A*^, and *Kirrel3* ^*Q128A/Q128A*^ mouse brains. A blinded analysis measuring glomerular size and number using Fiji software was performed on 8 to 10 consecutive sections from both AOBs that contained anterior and posterior regions of similar sizes at comparable medial-lateral level. Each VGLUT2 positive unit surrounded by a region of non-innervated neuropil was defined as a glomerulus and was manually traced on Fiji by an expert. The number of glomeruli and their sizes was measured from each section examined and the average values for each mouse brain was calculated. The anterior-posterior AOB border was identified by staining sections with BS lectin (data not shown). Littermates were used for analyses.

### Analysis of Kirrel3 protein expression

Whole brain lysates were harvested using 20 mM HEPES, 320 mM sucrose with protease inhibitors (0.1 µg/µl Leupeptin/Aprotinin and 1 mM PMSF). Protein concentration of lysate was determined using the Bio-Rad protein concentration assay and equal amounts of protein were loaded and subjected to SDS-PAGE gel electrophoresis followed by transfer to PVDF membranes (Immobilon-P). Membranes were probed with mouse anti-Kirrel3 1:100 (Neuromab Clone N321C/49; catalog no. 75-333) that detects the long and short isoforms of Kirrel3, rat anti-HSC70/HSP73 1:10,000 (Enzo Life Sciences), or rabbit anti-β-actin 1:1,000 (Cell Signalling Technology), followed with appropriate HRP-conjugated secondary antibodies. Blots were developed using SuperSignal West Femto Kit (Thermo Scientific). Quantification of relative protein levels was performed on scanned immunoblots using Fiji software.

### Cell surface biotinylation in acute brain slices

Cell surface biotinylation assay was performed as described (Gabriel et al., 2014). Acute mouse brain slices were prepared from *Kirrel3*^*Q128A/Q128A*^ and *Kirrel3*^+/+^ mice, placed in ACSF and allowed to recover in a recovery chamber for 1 hour at room temperature. Two slices were incubated in in chilled ACSF containing 1 mg/ml EZ-link Sulfo-NHS-SS-Biotin (Thermo Fisher Scientific) for 45 minutes. The biotin was then quenched with two washes of 10 mM Glycine in ACSF at 4°C followed by three washes in ice cold ACSF for 5 min. Slices were then harvested in RIPA buffer (10 mM Tris pH 7.45, 150 mM NaCl, 1 mM EDTA, 1% Triton X-100, 0.1% SDS and 1% sodium deoxycholate) supplemented with protease and phosphatase inhibitors (100 mM PMSF and PhosSTOP tablet (Roche)). Biotinylated proteins were precipitated with streptavidin-agarose beads (Thermo Fisher Scientific) at 4°C overnight, washed, and eluted from beads using SDS-PAGE reducing sample buffer. Equal amounts of protein were then subjected to a Western Blot analysis as described above.

## References

Afonine, P.V., Grosse-Kunstleve, R.W., Echols, N., Headd, J.J., Moriarty, N.W., Mustyakimov, M., Terwilliger, T.C., Urzhumtsev, A., Zwart, P.H., and Adams, P.D. (2012). Towards automated crystallographic structure refinement with phenix.refine. Acta Crystallogr. D Biol. Crystallogr. 68, 352–367.

Bear, D.M., Lassance, J.-M., Hoekstra, H.E., and Datta, S.R. (2016). The Evolving Neural and Genetic Architecture of Vertebrate Olfaction. Curr Biol 26, R1039–R1049.

Belluscio, L., Koentges, G., Axel, R., and Dulac, C. (1999). A map of pheromone receptor activation in the mammalian brain. Cell 97, 209–220.

Bhalla, K., Luo, Y., Buchan, T., Beachem, M.A., Guzauskas, G.F., Ladd, S., Bratcher, S.J., Schroer, R.J., Balsamo, J., DuPont, B.R., et al. (2008). Alterations in CDH15 and KIRREL3 in patients with mild to severe intellectual disability. Am. J. Hum. Genet. 83, 703–713.

Brautigam, C.A. (2015). Calculations and Publication-Quality Illustrations for Analytical Ultracentrifugation Data. Meth. Enzymol. 562, 109–133.

Brignall, A.C., and Cloutier, J.-F. (2015). Neural map formation and sensory coding in the vomeronasal system. Cell. Mol. Life Sci. 72, 4697–4709.

Brignall, A.C., Raja, R., Phen, A., Prince, J.E.A., Dumontier, E., and Cloutier, J.-F. (2018). Loss of Kirrel family members alters glomerular structure and synapse numbers in the accessory olfactory bulb. Brain Struct Funct 223, 307–319.

Bulchand, S., Menon, S.D., George, S.E., and Chia, W. (2010). The intracellular domain of Dumbfounded affects myoblast fusion efficiency and interacts with Rolling pebbles and Loner. PLoS One 5, e9374.

Cheng, S., Park, Y., Kurleto, J.D., Jeon, M., Zinn, K., Thornton, J.W., and Özkan, E. (2019). Family of neural wiring receptors in bilaterians defined by phylogenetic, biochemical, and structural evidence. Proc. Natl. Acad. Sci. U.S.A. 116, 9837–9842.

Chia, P.H., Chen, B., Li, P., Rosen, M.K., and Shen, K. (2014). Local F-actin Network Links Synapse Formation and Axon Branching. Cell 156, 208–220.

Darriba, D., Posada, D., Kozlov, A.M., Stamatakis, A., Morel, B., and Flouri, T. (2020). ModelTest-NG: A New and Scalable Tool for the Selection of DNA and Protein Evolutionary Models. Mol. Biol. Evol. 37, 291–294.

De Rubeis, S., He, X., Goldberg, A.P., Poultney, C.S., Samocha, K., Cicek, A.E., Kou, Y., Liu, L., Fromer, M., Walker, S., et al. (2014). Synaptic, transcriptional and chromatin genes disrupted in autism. Nature 515, 209–215.

Dehal, P., and Boore, J.L. (2005). Two rounds of whole genome duplication in the ancestral vertebrate. PLoS Biol. 3, e314.

Del Punta, K., Puche, A., Adams, N.C., Rodriguez, I., and Mombaerts, P. (2002). A divergent pattern of sensory axonal projections is rendered convergent by second-order neurons in the accessory olfactory bulb. Neuron 35, 1057–1066.

Donoviel, D.B., Freed, D.D., Vogel, H., Potter, D.G., Hawkins, E., Barrish, J.P., Mathur, B.N., Turner, C.A., Geske, R., Montgomery, C.A., et al. (2001). Proteinuria and perinatal lethality in mice lacking NEPH1, a novel protein with homology to NEPHRIN. Mol Cell Biol 21, 4829–4836.

Dulac, C., and Torello, A.T. (2003). Molecular detection of pheromone signals in mammals: from genes to behaviour. Nat Rev Neurosci 4, 551–562.

Edgar, R.C. (2004). MUSCLE: multiple sequence alignment with high accuracy and high throughput. Nucleic Acids Res. 32, 1792–1797.

Emsley, P., Lohkamp, B., Scott, W.G., and Cowtan, K. (2010). Features and development of Coot. Acta Crystallogr. D Biol. Crystallogr. 66, 486–501.

Franke, D., and Svergun, D.I. (2009). DAMMIF, a program for rapid ab-initio shape determination in small-angle scattering. J Appl Crystallogr 42, 342–346.

Gabriel, L.R., Wu, S., and Melikian, H.E. (2014). Brain slice biotinylation: an ex vivo approach to measure region-specific plasma membrane protein trafficking in adult neurons. J Vis Exp e51240.

Gerke, P., Huber, T.B., Sellin, L., Benzing, T., and Walz, G. (2003). Homodimerization and heterodimerization of the glomerular podocyte proteins nephrin and NEPH1. J Am Soc Nephrol 14, 918–926.

Gerke, P., Sellin, L., Kretz, O., Petraschka, D., Zentgraf, H., Benzing, T., and Walz, G. (2005). NEPH2 is located at the glomerular slit diaphragm, interacts with nephrin and is cleaved from podocytes by metalloproteinases. J Am Soc Nephrol 16, 1693–1702.

Gerke, P., Benzing, T., Höhne, M., Kispert, A., Frotscher, M., Walz, G., and Kretz, O. (2006). Neuronal expression and interaction with the synaptic protein CASK suggest a role for Neph1 and Neph2 in synaptogenesis. J. Comp. Neurol. 498, 466–475.

Harita, Y., Kurihara, H., Kosako, H., Tezuka, T., Sekine, T., Igarashi, T., and Hattori, S. (2008). Neph1, a component of the kidney slit diaphragm, is tyrosine-phosphorylated by the Src family tyrosine kinase and modulates intracellular signaling by binding to Grb2. J. Biol. Chem. 283, 9177–9186.

Hisaoka, T., Komori, T., Kitamura, T., and Morikawa, Y. (2018). Abnormal behaviours relevant to neurodevelopmental disorders in Kirrel3-knockout mice. Sci Rep 8, 1408.

Hopkins, J.B., Gillilan, R.E., and Skou, S. (2017). BioXTAS RAW: improvements to a free open-source program for small-angle X-ray scattering data reduction and analysis. J Appl Crystallogr 50, 1545–1553.

Huber, T.B., Schmidts, M., Gerke, P., Schermer, B., Zahn, A., Hartleben, B., Sellin, L., Walz, G., and Benzing, T. (2003). The carboxyl terminus of Neph family members binds to the PDZ domain protein zonula occludens-1. J Biol Chem 278, 13417–13421.

Kozlov, A.M., Darriba, D., Flouri, T., Morel, B., and Stamatakis, A. (2019). RAxML-NG: a fast, scalable and user-friendly tool for maximum likelihood phylogenetic inference. Bioinformatics 35, 4453–4455.

Laue, T.M., Shah, B.D., Ridgeway, R.M., and Pelletier, S.L. (1992). Computer-aided interpretation of analytical sedimentation data for proteins. In Analytical Ultracentrifugation in Biochemistry and Polymer Science, S.E. Harding, A.J. Rowe, and J.C. Horton, eds. (Cambridge, UK: The Royal Society of Chemistry), pp. 90–125.

Liebschner, D., Afonine, P.V., Baker, M.L., Bunkóczi, G., Chen, V.B., Croll, T.I., Hintze, B., Hung, L.W., Jain, S., McCoy, A.J., et al. (2019). Macromolecular structure determination using X-rays, neutrons and electrons: recent developments in Phenix. Acta Crystallogr D Struct Biol 75, 861–877.

Manalastas-Cantos, K., Konarev, P.V., Hajizadeh, N.R., Kikhney, A.G., Petoukhov, M.V., Molodenskiy, D.S., Panjkovich, A., Mertens, H.D.T., Gruzinov, A., Borges, C., et al. (2021). ATSAS 3.0: expanded functionality and new tools for small-angle scattering data analysis. J Appl Cryst 54, 343–355.

Martin, E.A., Muralidhar, S., Wang, Z., Cervantes, D.C., Basu, R., Taylor, M.R., Hunter, J., Cutforth, T., Wilke, S.A., Ghosh, A., et al. (2015). The intellectual disability gene Kirrel3 regulates target-specific mossy fiber synapse development in the hippocampus. Elife 4, e09395.

McCoy, A.J., Grosse-Kunstleve, R.W., Adams, P.D., Winn, M.D., Storoni, L.C., and Read, R.J. (2007). Phaser crystallographic software. J Appl Crystallogr 40, 658–674.

Mombaerts, P., Wang, F., Dulac, C., Chao, S.K., Nemes, A., Mendelsohn, M., Edmondson, J., and Axel, R. (1996). Visualizing an olfactory sensory map. Cell 87, 675–686.

Nakashima, A., Ihara, N., Shigeta, M., Kiyonari, H., Ikegaya, Y., and Takeuchi, H. (2019). Structured spike series specify gene expression patterns for olfactory circuit formation. Science 365.

Otwinowski, Z., and Minor, W. (1997). [20] Processing of X-ray diffraction data collected in oscillation mode. In Macromolecular Crystallography, Part A, C.W. Carter Jr., and R.M. Sweet, eds. (Academic Press (New York), pp. 307–326.

Özkan, E., Carrillo, R.A., Eastman, C.L., Weiszmann, R., Waghray, D., Johnson, K.G., Zinn, K., Celniker, S.E., and Garcia, K.C. (2013). An Extracellular Interactome of Immunoglobulin and LRR Proteins Reveals Receptor-Ligand Networks. Cell 154, 228–239.

Özkan, E., Chia, P.H., Wang, R.R., Goriatcheva, N., Borek, D., Otwinowski, Z., Walz, T., Shen, K., and Garcia, K.C. (2014). Extracellular Architecture of the SYG-1/SYG-2 Adhesion Complex Instructs Synaptogenesis. Cell 156, 482–494.

Petoukhov, M.V., and Svergun, D.I. (2005). Global rigid body modeling of macromolecular complexes against small-angle scattering data. Biophys J 89, 1237–1250.

Poncelet, G., and Shimeld, S.M. (2020). The evolutionary origins of the vertebrate olfactory system. Open Biol 10, 200330.

Prince, J.E.A., Brignall, A.C., Cutforth, T., Shen, K., and Cloutier, J.-F. (2013). Kirrel3 is required for the coalescence of vomeronasal sensory neuron axons into glomeruli and for male-male aggression. Development 140, 2398–2408.

Ramos, R.G., Igloi, G.L., Lichte, B., Baumann, U., Maier, D., Schneider, T., Brandstätter, J.H., Fröhlich, A., and Fischbach, K.F. (1993). The irregular chiasm C-roughest locus of Drosophila, which affects axonal projections and programmed cell death, encodes a novel immunoglobulin-like protein. Genes Dev 7, 2533–2547.

Rodriguez, I., Feinstein, P., and Mombaerts, P. (1999). Variable patterns of axonal projections of sensory neurons in the mouse vomeronasal system. Cell 97, 199–208.

Roy, A., Kucukural, A., and Zhang, Y. (2010). I-TASSER: a unified platform for automated protein structure and function prediction. Nat Protoc 5, 725–738.

Ruiz-Gómez, M., Coutts, N., Price, A., Taylor, M.V., and Bate, M. (2000). Drosophila dumbfounded: a myoblast attractant essential for fusion. Cell 102, 189–198.

Sakurai, T., Watanabe, S., Kamiyoshi, A., Sato, M., and Shindo, T. (2014). A single blastocyst assay optimized for detecting CRISPR/Cas9 system-induced indel mutations in mice. BMC Biotechnol 14, 69.

Sanes, J.R., and Zipursky, S.L. (2020). Synaptic Specificity, Recognition Molecules, and Assembly of Neural Circuits. Cell 181, 536–556.

Schuck, P. (2000). Size-distribution analysis of macromolecules by sedimentation velocity ultracentrifugation and Lamm equation modeling. Biophys. J. 78, 1606–1619.

Schuck, P. (2003). On the analysis of protein self-association by sedimentation velocity analytical ultracentrifugation. Anal Biochem 320, 104–124.

Sellin, L., Huber, T.B., Gerke, P., Quack, I., Pavenstädt, H., and Walz, G. (2003). NEPH1 defines a novel family of podocin interacting proteins. FASEB J 17, 115–117.

Serizawa, S., Miyamichi, K., Takeuchi, H., Yamagishi, Y., Suzuki, M., and Sakano, H. (2006). A neuronal identity code for the odorant receptor-specific and activity-dependent axon sorting. Cell 127, 1057–1069.

Shen, K., and Bargmann, C.I. (2003). The immunoglobulin superfamily protein SYG-1 determines the location of specific synapses in C. elegans. Cell 112, 619–630.

Svergun, D.I. (1992). Determination of the regularization parameter in indirect-transform methods using perceptual criteria. J Appl Crystallogr 25, 495–503.

Taylor, M.R., Martin, E.A., Sinnen, B., Trilokekar, R., Ranza, E., Antonarakis, S.E., and Williams, M.E. (2020). Kirrel3-Mediated Synapse Formation Is Attenuated by Disease-Associated Missense Variants. J. Neurosci. 40, 5376–5388.

Tria, G., Mertens, H.D.T., Kachala, M., and Svergun, D.I. (2015). Advanced ensemble modelling of flexible macromolecules using X-ray solution scattering. IUCrJ 2, 207–217.

Vaddadi, N., Iversen, K., Raja, R., Phen, A., Brignall, A., Dumontier, E., and Cloutier, J.-F. (2019). Kirrel2 is differentially required in populations of olfactory sensory neurons for the targeting of axons in the olfactory bulb. Development 146.

Völker, L.A., Maar, B.A., Pulido Guevara, B.A., Bilkei-Gorzo, A., Zimmer, A., Brönneke, H., Dafinger, C., Bertsch, S., Wagener, J.-R., Schweizer, H., et al. (2018). Neph2/Kirrel3 regulates sensory input, motor coordination, and home-cage activity in rodents. Genes Brain Behav 17, e12516.

Wolff, T., and Ready, D.F. (1991). Cell death in normal and rough eye mutants of Drosophila. Development 113, 825–839.

Yang, Z. (2007). PAML 4: phylogenetic analysis by maximum likelihood. Mol. Biol. Evol. 24, 1586–1591.

Yang, J., and Zhang, Y. (2015). I-TASSER server: new development for protein structure and function predictions. Nucleic Acids Res 43, W174–181.

Yesildag, B., Bock, T., Herrmanns, K., Wollscheid, B., and Stoffel, M. (2015). Kin of IRRE-like Protein 2 Is a Phosphorylated Glycoprotein That Regulates Basal Insulin Secretion. J Biol Chem 290, 25891–25906.

