## Supplemental Figures for "Molecular and Structural Basis of Olfactory Sensory Neuron Coalescence by Kirrel Receptors"

### SUPPLEMENTAL FIGURE LEGENDS

**Figure S1.** Structural comparison of mouse Kirrel3 D1 homodimer with orthologous *Drosophila* Kirre (Duf) D1 homodimer (**A**), *Drosophila* Rst D1 homodimer (**B**), and *C. elegans* SYG-1-SYG-2 D1 heterodimer (**C**). D1 subunits on the left hand side in all panels have been superimposed to highlight differences in exact dimerization topologies.

**Figure S2. A,B.** Hydrogen bonding networks conserved between Kirrel2 (**A**) and Kirrel3 (**B**) homodimerization interfaces. **C.** The conserved networks are formed by a Gln and an Arg in F and G strands respectively, and by main-chain atoms in a conserved stretch in the CD loop.

**Figure S3.** Ancestral vertebrate and gnathostome Kirrel D1 sequences in a multiple sequence alignment with mouse Kirrel3 D1.

**Figure S4. A,B.** Size-exclusion chromatography profiles for mouse Kirrel2 D1 mutants loaded at lower (A) and higher concentrations (B). **C,D.** Size-exclusion chromatography profiles for mouse Kirrel3 D1+D2 mutants loaded at lower (C) and higher concentrations (D). **E,F.** Size-exclusion chromatography profiles for mKirrel2 and mKirrel3 WT and mutants loaded at various quantities. The coloring scheme follows that in (B) for mKirrel2 variants (E), and that in (D) for mKirrel3 variants. **G,H.** Binding isotherms for mKirrel2 and mKirrel3 ectodomains plotted as Kirrel concentration against sedimentation coefficients.

**Figure S5. A,B.** Pair distance distribution, $P(r)$, (A) and dimensionless Kratky plot for mKirrel2 WT ectodomain (B). Blue vertical lines in (B) are measurement errors. Dashed red lines show predicted plots for a rigid, globular molecule with the same $R_g$. **C.** Differential refractive index and molar mass measurements for Kirrel ectodomains. See Table S3 for experimental details. **D.** Bead model from *DAMMIF* for mKirrel3 Q128A ectodomain overlayed with the highest prevalence *EOM* model. **E.** Bead model from *DAMMIF* for mKirrel3 WT ectodomain overlayed with the *SASREF* model.

**Figure S6.** Cell surface display of FLAG-tagged mKirrel3, WT (left) and Q128A mutant (right), on HEK293 cells.

**Table S1, related to Figure 2.** Data and refinement statistics for x-ray crystallography.

**Table S2, related to Figure 5.** Data collection details and analysis statistics for SAXS experiments.

**Table S3, related to Figure 5.** Data collection details and analysis statistics for MALS experiments.

**Source Data 1.** Phylogenetic tree with bootstrap support values for nodes in Newick format. Abbreviations for species used: (Mammals) Mmus, *Mus musculus*; Hsap, *Homo sapiens*; Clup, *Canis lupus familiaris*; Lafr, *Loxodonta africana*; Mdom, *Monodelphis domestica*; Oana, *Ornithorhynchus anatinus*; (Aves) Ggal, *Gallus gallus*; Nper, *Nothoprocta perdicaria*; (Reptilia) Acar, *Anolis carolinensis*; Cpic, *Chrysemys picta bellii*; (Amphibia) Xtro, *Xenopus tropicalis*; Npar, *Nanorana parkeri*; Muni, *Microcaecilia unicolor*; Gser, *Geotrypetes seraphini*; (Sarcopterygii) Lcha, *Latimeria chalumnae*; (Actinoptergyii); Ecal, *Erpetoichthys calabaricus*; Arut, *Acipenser ruthenus*; Locu, *Lepisosteus oculatus*; Drer, *Danio rerio*; (Chondrichthyes) Cmil, *Callorhinchus milii*; Arad, *Amblyraja radiata*; Stor, *Scyliorhinus torazame*; Cpun, *Chiloscyllium punctatum*;

(Cyclostomata) Pmar, *Petromyzon marinus*; Ebur, *Eptatretus burgeri*; Lcam, *Lethenteron camtschaticum*; (Urochordata) Cint, *Ciona intestinalis*; Pmam, *Phallusia mammillata*; (Cephalochordata) Bbel, *Branchiostoma belcheri*; (Ambulacraria) Spur, *Strongylocentrotus purpuratus*; Skow, *Saccoglossus kowalevskii*; (Xenacoelomorpha) Ipul, *Isodiametra pulchra*; Msti, *Meara stichopi*; (Spiralia) Hmic, *Hymenolepis microstoma*; Ctel, *Capitella teleta*; Lana, *Lingula anatina*; Obim, *Octopus bimaculoides*; Cvir, *Crassostrea virginica*; (Ecdysozoa) Pcau, *Priapulus caudatus*; Dmel, *Drosophila melanogaster*; Tcas, *Tribolium castaneum*; Isca, *Ixodes scapularis*; Cscu, *Centruroides sculpturatus*; Hduj, *Hypsibius dujardini*; Rvar, *Ramazzotius varieornatus*; Cele, *Caenorhabditis elegans*; Bmal, *Brugia malayi*; Tspi, *Trichinella spiralis*; Sbat, *Soboliphyme baturini*.

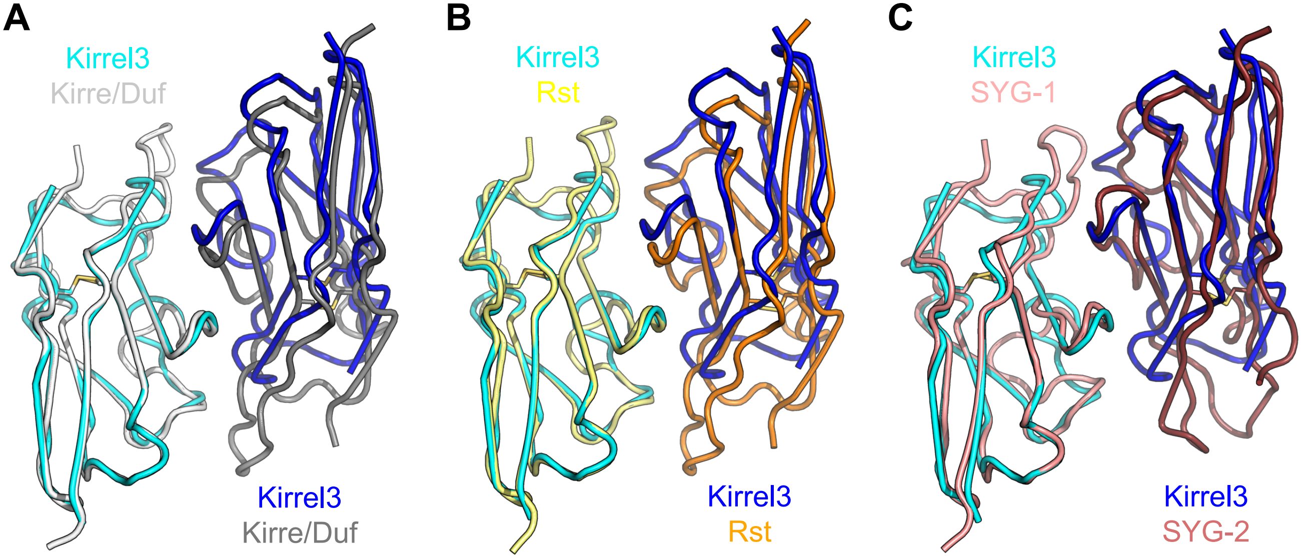


**Figure S1.** Structural comparison of mouse Kirrel3 D1 homodimer with orthologous *Drosophila* Kirre (Duf) D1 homodimer **(A)**, *Drosophila* Rst D1 homodimer **(B)**, and *C. elegans* SYG-1-SYG-2 D1 heterodimer **(C)**. D1 subunits on the left hand side in all panels have been superimposed to highlight differences in exact dimerization topologies.

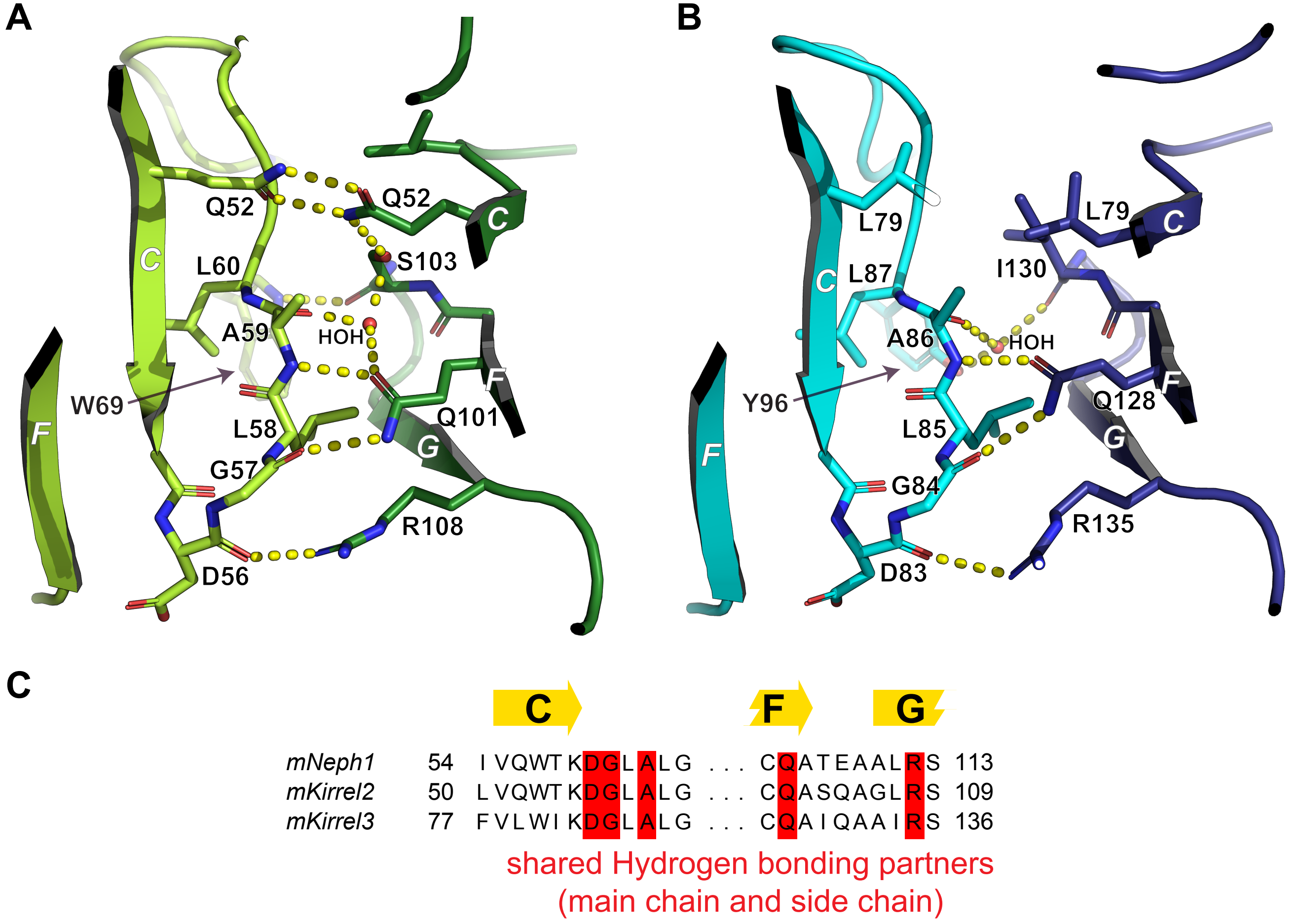


**Figure S2. A,B.** Hydrogen bonding networks conserved between Kirrel2 (A) and Kirrel3 (B) homodimerization interfaces. **C.** The conserved networks are formed by a Gln and an Arg in F and G strands respectively, and by main-chain atoms in a conserved stretch in the CD loop.

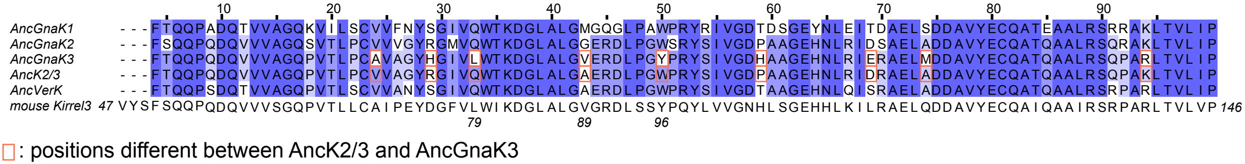

Figure S3. Ancestral vertebrate and gnathostome Kirrel D1 sequences in a multiple sequence alignment with mouse Kirrel3 D1.

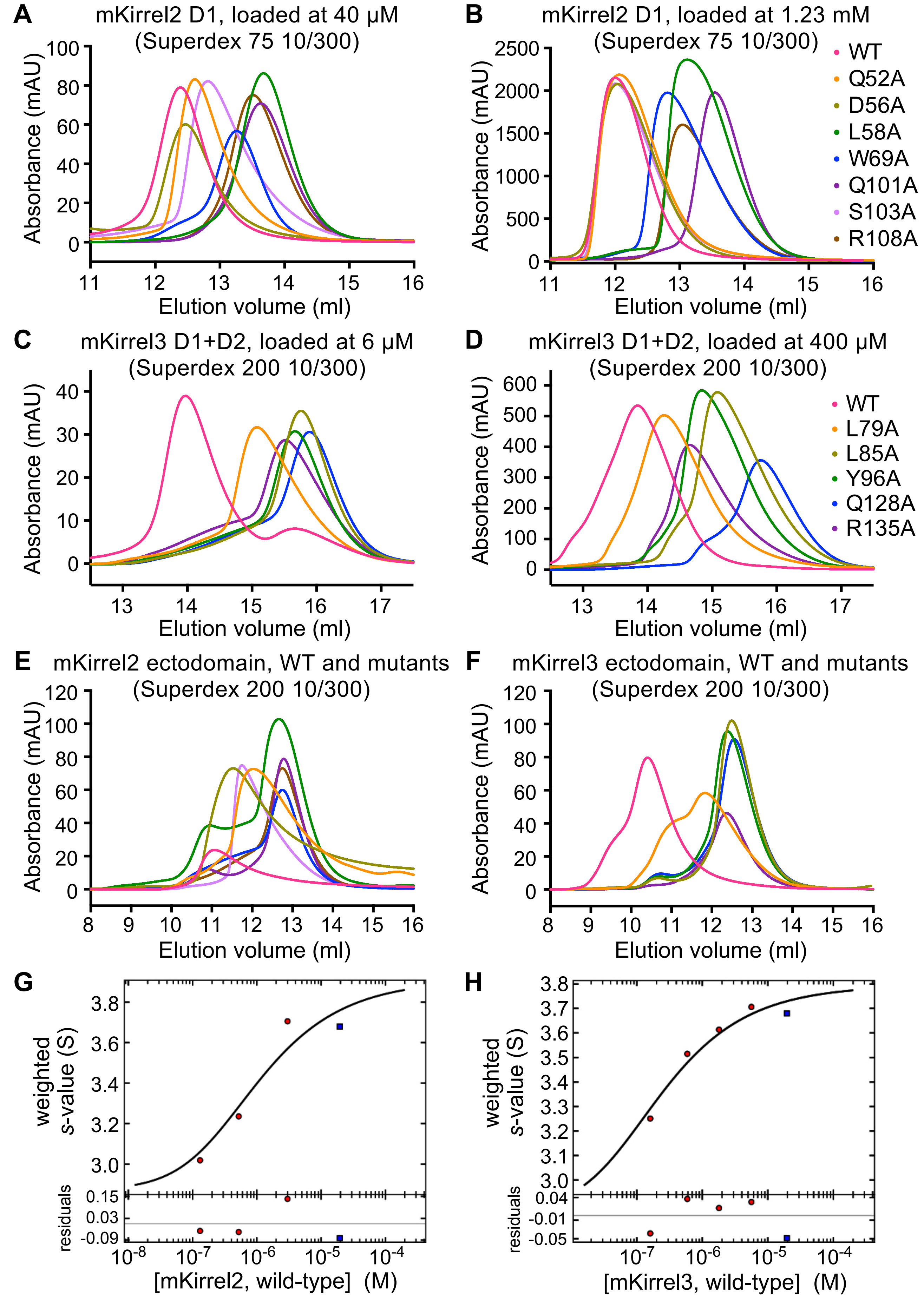


**Figure S4. A,B.** Size-exclusion chromatography profiles for mouse Kirrel2 D1 mutants loaded at lower (A) and higher concentrations (B). **C,D.** Size-exclusion chromatography profiles for mouse Kirrel3 D1+D2 mutants loaded at lower (C) and higher concentrations (D). **E,F.** Size-exclusion chromatography profiles for mKirrel2 and mKirrel3 WT and mutants loaded at various quantities. The coloring scheme follows that in (B) for mKirrel2 variants (E), and that in (D) for mKirrel3 variants. **G,H.** Binding isotherms for mKirrel2 and mKirrel3 ectodomains plotted as Kirrel concentration against sedimentation coefficients.

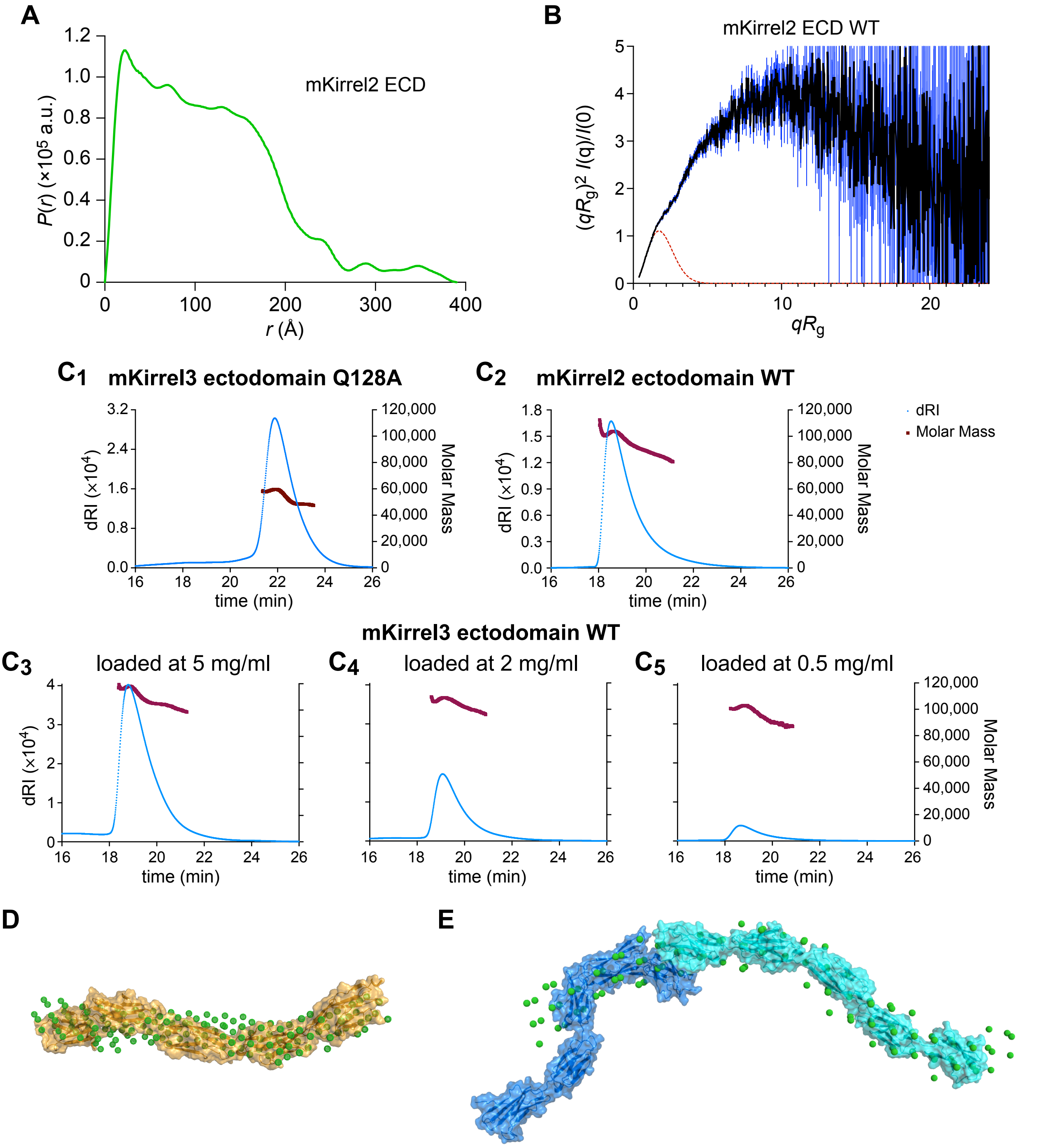


**Figure S5. A,B.** Pair distance distribution, $P(r)$ ,(A) and dimensionless Kratky plot for mKirrel2 WT ectodomain (B). Blue vertical lines in (B) are measurement errors. Dashed red lines show predicted plots for a rigid, globular molecule with the same $R_g$. **C.** Differential refractive index and molar mass measurements for Kirrel ectodomains. See Table S3 for experimental details. **D.** Bead model from *DAMMIF* for mKirrel3 Q128A ectodomain overlayed with the highest prevalence *EOM* model. **E.** Bead model from *DAMMIF* for mKirrel3 WT ectodomain overlayed with the *SASREF* model.

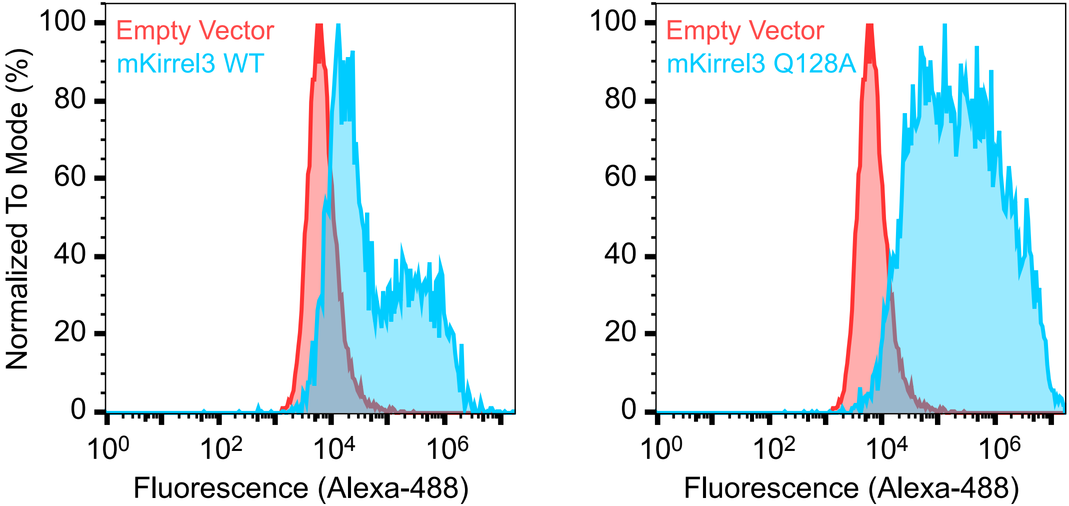


**Figure S6.** Cell surface display of FLAG-tagged mKirrel3, WT (left) and Q128A mutant (right), on HEK293 cells.

**Table S1, *related to Figure 2.*** Data and refinement statistics for x-ray crystallography.

| | Mouse Kirrel2 D1 | Mouse Kirrel3 D1 |
|---|---|---|
| **Data Collection** | | |
| Beamline | SSRL 12-2 | APS 24-ID-E |
| Space Group | *C*2 | $P6_3$ |
| *Cell Dimensions* | | |
| $a, b, c$ (Å) | 109.14, 48.44, 44.10 | 63.24, 63.24, 347.81 |
| $\alpha, \beta, \gamma$ (°) | 90, 107.01, 90 | 90, 90, 120 |
| Twin law (Fraction) | – | -k,-h,-l (0.31)[†] |
| Resolution (Å) | 50-1.80 (1.83-1.80)* | 50-1.95 (1.98-1.95) |
| $R_{sym}$ (%) | 5.6 (17.3) | 11.4 (85.3) |
| $\langle I\rangle/\langle\sigma(I)\rangle$ | 21.9 (5.3) | 14.6 (1.9) |
| $CC_{1/2}$ in the highest resolution bin | 0.945 | 0.617 |
| Completeness (%) | 94.2 (69.1) | 99.9 (99.8) |
| Redundancy | 3.3 (2.6) | 4.4 (4.4) |
| **Refinement** | | |
| Resolution (Å) | 50-1.80 (1.90-1.80)* | 50.1-1.95 (1.98-1.95) |
| Reflections | 19,361 | 56,984 |
| $R_{cryst}$ (%) | 15.49 (19.50) | 16.02 (22.95) |
| $R_{free}$ (%)** | 18.09 (25.18) | 19.53 (27.48) |
| *Number of atoms* | | |
| Protein | 1633 | 6408 |
| Ligand | 2 | 8 |
| Water | 337 | 336 |
| *Average B-factors* ($Å^2$) | | |
| All | 20.1 | 38.1 |
| Protein | 18.1 | 38.5 |
| Ligand/Glycans | 11.8 | 44.9 |
| Solvent | 30.0 | 31.4 |
| *R.m.s. deviations from ideality* | | |
| Bond Lengths (Å) | 0.004 | 0.005 |
| Bond Angles (°) | 0.632 | 0.684 |
| *Ramachandran plot* | | |
| Favored (%) | 98.52 | 96.19 |
| Outliers (%) | 0.00 | 0.00 |
| Rotamer Outliers (%) | 0.00 | 0.14 |
| All-atom Clashscore[†] | 0.00 | 3.43 |

* The values in parentheses are for reflections in the highest resolution bin.
** 5% of reflections (971 for Kirrel2 and 2846 for Kirrel3) were not used during refinement for cross validation.
[†] Clashscores and twin fraction were calculated by phenix.refine (*Phenix* version 1.19.1-4122).

**Table S2**, ***related to Figure 5.*** Data collection details and analysis statistics for SAXS experiments.

| | mKirrel2 ECD | mKirrel3 ECD | mKirrel3 ECD Q128A |
|---|---|---|---|
| *Data Collection and Sample Details* | | | |
| Experiment setup | SEC-SAXS-MALS | | |
| Instrument | BioCAT facility at APS beamline 18-ID with Pilatus3 X 1M (Dectris) detector | | |
| Wavelength (Å) | 1.033 | | |
| Camera length (m) | 3.655 | | |
| $q$-measurement range (Å$^{-1}$) | 0.0044 to 0.35 | | |
| Exposure time (s) | 0.5 | | |
| Exposure period (s) | 1.0 | | |
| Flow rate (ml/min) | 0.6 | | |
| Chromatography column | Superdex 200 10/300 Increase | | |
| Buffer | 10 mM HEPES pH 7.2, 150 mM NaCl | | |
| Temperature | 22°C | | |
| Software | BioXTAS RAW version 2.0.2 | | |
| Loading concentration (mg/ml) | 3.7 | 2.0 | 5.0 |
| Loading volume (µl) | 425 | | |
| *Structural Parameters* | | | |
| Guinier Analysis* | | | |
| $I(0)$ (cm$^{-1}$) | 0.0231 ± 0.000186 | 0.0596 ± 0.000169 | 0.0247 ± 0.000082 |
| $R_g$ (Å) | 88.67 ± 1.28 | 93.17 ± 0.49 | 54.42 ± 0.30 |
| $q$ range (Å$^{-1}$) | 0.005 to 0.0133 | 0.0047 to 0.0124 | 0.005 to 0.0227 |
| $q_{min}R_g$ to $q_{max}R_g$ | 0.444 to 1.178 | 0.439 to 1.158 | 0.272 to 1.237 |
| Coefficient of Correlation, $R^2$ | 0.981 | 0.996 | 0.991 |
| $P(r)$ Analysis (from GNOM**) | | | |
| $I(0)$ (cm$^{-1}$) | 0.0235 ± 0.000234 | 0.0595 ± 0.000155 | 0.0251 ± 0.000085 |
| $R_g$ (Å) | 95.97 ± 2.21 | 95.59 ± 0.40 | 58.69 ± 0.31 |
| $D_{max}$ (Å) | 390 | 345 | 213 |
| $q$ range (Å$^{-1}$) | 0.0050 to 0.3497 | 0.0047 to 0.3497 | 0.0050 to 0.3497 |
| $\chi^2$ | 0.856 | 1.050 | 1.104 |
| Porod Volume estimate (Å$^3$) | 152000 | 157000 | 109000 |
| Molecular weight based on volume of correlation ($V_c$) | 80,800 | 94,600 | 45,400 |

* Performed in BioXTAS RAW (Hopkins, J.B. *et al.* (2017) *J Appl Crystallogr* **50**, 1545-1553).
** Svergun, D.I. (1992). *J Appl Crystallogr* **25**, 495-503.

**Table S3, *related to Figure 5*.** Data collection details and analysis statistics for MALS experiments.

| | mKirrel2 ECD | mKirrel3 ECD Q128A | mKirrel3 ECD WT run 1 | mKirrel3 ECD WT run 2 | mKirrel3 ECD WT run 3 |
|---|---|---|---|---|---|
| Experiment Type | SEC-SAXS-MALS | | | | |
| Instrument | DAWN HELEOS with Optilab T-rEX at BioCAT (APS beamline 18-ID) | | | | |
| Wavelength (nm) | 660 (for light scattering) and 658 (for RI) | | | | |
| Flow rate (ml/min) | 0.6 | | | | |
| Chromatography column | Superdex 200 10/300 Increase | | | | |
| Buffer | 10 mM HEPES pH 7.2, 150 mM NaCl | | | | |
| Refractive index of the solvent (assumed) | 1.331 | | | | |
| Temperature | 25°C | | | | |
| Software | ASTRA version 7.3.2.19 (Wyatt) | | | | |
| Loading volume (µl) | 425 | | | | |
| Loading concentration (mg/ml) | 3.7 | 2.0 | 5.0 | 2.0 | 0.5 |
| *dn*/*dc** | 0.182 | 0.181 | 0.181 | 0.181 | 0.181 |
| $M_n$** | 97,200 (±6.2%) | 54,830 (±7.5%) | 111,100 (±6.5%) | 105,400 (±4.6%) | 97,910 (±5.2%) |
| $M_w$† | 97,630 (±6.2%) | 55,200 (±7.3%) | 111,400 (±6.5%) | 105,500 (±4.6%) | 98,170 (±5.2%) |

* *dn*/*dc* values are estimated based on predicted N-linked glycosylation content.
** Number-averaged molar mass.
† Weight-averaged molar mass.
